# Extreme long-range migration distorts flower colour clines in an *Antirrhinum* hybrid zone

**DOI:** 10.1101/2025.06.25.661303

**Authors:** Parvathy Surendranadh, Sean Stankowski, David L. Field, Nicholas H. Barton

## Abstract

Hybrid zones are natural laboratories for the study of speciation, allowing researchers to quantify selection, dispersal, and barriers to gene flow in the real world. We analyzed a flower-colour hybrid zone between two varieties of the common snapdragon, *Antirrhinum majus* subspecies *majus*, based on a dataset of 22,494 individuals sampled over 10 consecutive years (Field et al. in prep). We found narrow geographic clines at six SNP markers that are tightly linked to known flower colour loci, with widths ranging from 0.8 to 5.5 km. The clines have a ‘stepped’ shape, which is expected when strong linkage disequilibrium generated by diffusive dispersal into the hybrid zone, increases the effective selection at the cline centre. However, the observed LD (mean 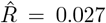) was far too weak to explain the stepped cline shapes. Instead, we show that the stepped clines are shaped primarily by the observed long-distance dispersal. We used a novel simulation framework that accounts for realistic dispersal and multilocus selection to estimate the direct selection (1.5% to 18.5%) and indirect selection due to LD (1.2% to 2.4%) on each colour locus. Finally, we show that the barrier to the flow of neutral alleles through diffusive dispersal causes negligible distortion to the clines, while the reduction in gene flow due to selection on long-range migrants is substantial (∼50%). The results shed new light onto the processes that shape hybrid zones, and highlight the need to account for the effects of realistic dispersal in theoretical and empirical studies of hybrid zones.

## 1 Introduction

Hybrid zones are narrow areas where genetically differentiated populations come into contact with one another, leading to the production of hybrid offspring (Barton and Hewitt, 1985; Hewitt, 1988). They are found in a wide range of organisms and environments, and are considered natural laboratories for studying divergence and speciation in the wild (Hewitt, 1988; Harrison, 1993). Most well-studied hybrid zones are thought to reflect the long-term balance between selection, which keeps them narrow, and dispersal, which acts to widen the zone (Barton and Hewitt, 1985). They have been extensively studied theoretically, providing a framework for inferring evolutionary processes from empirical data (Haldane, 1948; Slatkin, 1973; Endler, 1977; Barton, 1979a, 1983). Indeed, some of the most detailed inferences of selection, dispersal and barriers to gene flow have come from empirical studies of hybrid zones across diverse organisms, including fire-bellied toads (*Bombina* (e.g. Szymura and Barton (1986)), *Heliconius* butterflies (Mallet and Barton, 1989), *Carlia* lizards (Phillips et al., 2004), *Morabine* grasshoppers (Kawakami et al., 2009), and house mouse (Searle, 1991), to name a few.

Most inferences from hybrid zones are based on the properties of geographic clines (i.e., gradients in trait means or allele frequencies). When a single locus is under direct selection, and genes diffuse through the habitat, its corresponding allele frequency cline has a sigmoid shape (Fig. 1A), with width, *w*, proportional to the ratio between dispersal and selection. Thus, narrower clines imply stronger selection (Haldane, 1948; Bazykin, 1969; Slatkin, 1973). However, when clines at multiple selected loci coincide in a hybrid zone, they may have a ‘stepped’ shape in which a steep sigmoid gradient is flanked by shallow tails of introgression (Fig. 1B, Fig. S1, Barton (1983); Szymura and Barton (1986)). When multiple loci influence the trait under selection, each locus experiences an increase in effective selection due to positive associations (i.e., linkage disequilibrium; LD) between them at the cline centre (Barton, 1983; Kruuk et al., 1999). Specifically, the total selection experienced by a given locus is determined not only by its own direct effect on fitness, but also by indirect selection, i.e by its associations with other selected loci (even if unlinked). This effect is strongest in the centre of the cline, where LD is strongest, generating a barrier to the flow of alleles from one side of the hybrid zone to the other. However, as alleles escape their associations by recombining onto the alternative genetic background, the strength of indirect selection decreases and loci behave independently (i.e. weaker LD), leading to shallower tails of introgression away from the centre of the zone (Barton, 1983). The stepped cline model has been applied to a variety of empirical systems, enabling researchers to quantify a range of biological parameters, including the selection acting on each locus (*s*), the strength of the barrier to gene flow in units of geographic distance (*B*), and rates of introgression into the tails (*θ*)(e.g., (Szymura and Barton, 1986; Kawakami et al., 2009; Raufaste et al., 2005; Porter et al., 1997; Macholán et al., 2007). However, other factors can cause clines to have a stepped shape, so this need not always indicate a genetic barrier to gene flow.

**Figure 1.**
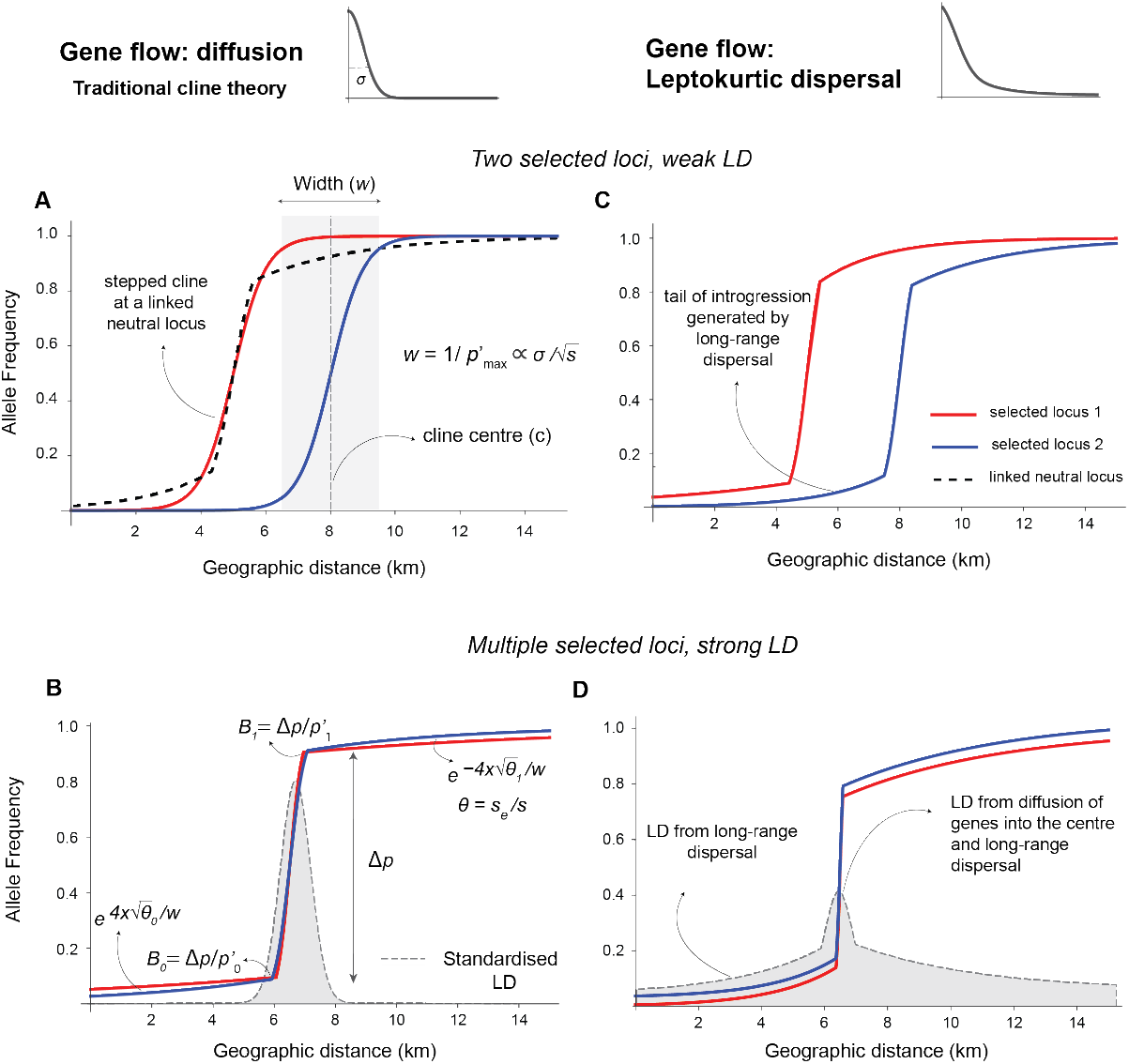
Illustration of clines under diffusive (left column) vs leptokurtic dispersal (right column). The red and blue curves in panels A-D show the cline shapes for 2 selected loci. Panels A and B show expected cline shapes under diffusive gene flow when LD is (A) weak and (B) strong. When LD is weak, clines have a sigmoid shape. The cline width (w) is defined as the inverse of the maximum gradient 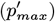 and is proportional to 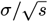, where *σ*is the standard deviation of the parent-offspring distance and *s* is the effective selection at the centre (*c*). When LD among selected loci is strong, it increases the effective selection at the centre, causing a ‘stepped’ cline with a steep central step flanked by shallower tails of introgression (see also Fig. S1). The tails take the form 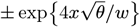, where *θ* is the ratio between effective selection at the centre (*s*) and selection against introgressing alleles (*s*_*e*_) away from the centre, and may differ to the left (*θ*_0_) and right (*θ*_1_). Under this scenario, the step reflects a genetic barrier to gene flow (*B*), described as *B* = Δ*p/p*^*′*^, where Δ*p* is the difference in allele frequency across the central step and *p*^*′*^ is the gradient of allele frequency at the edge of the step and the tail. (Panel C) When dispersal is leptokurtic, cline shapes may be stepped even if LD is weak. This is because long distance migration transports foreign alleles to the opposite tail in a single dispersal event, in contrast to diffusion, where alleles must cross the barrier via many small steps. In this scenario, the step is caused only by long-range dispersal, not LD. (Panel D) Both leptokurtic dispersal and LD contribute to the step. This scenario is distinguished from panel B by the presence of correlations between loci (i.e., standardised LD) in the tails of the cline which is generated by long-distance migration (grey distributions in B and D). Under diffusive gene flow, these correlations are only expected at the cline centre (as in panel B). In all four scenarios neutral loci linked to selected loci are expected to show stepped clines (dashed black curve in panel A), indicating a barrier to the flow of neutral alleles.

The pattern of dispersal may also influence cline shape. A key assumption of ‘traditional’ cline theory (Fig. 1A, B) is that the distribution of dispersal distance has finite moments, and that selection is weak enough that dispersal can be approximated as diffusion (Nagylaki, 1976; Slatkin, 1973). This enables dispersal to be described by a single parameter *σ*– the root mean square of parent-offspring distance along some axis, which can be estimated from mark-recapture studies or from LD at the cline centre (Mallet and Barton, 1989; Barton and Gale, 1993; Szymura and Barton, 1986, 1991). In reality, many species show leptokurtic dispersal, meaning that dispersal mostly occurs over short distances, but occasionally includes long-range movements (Nathan et al., 2008; Cayuela et al., 2018). These long-distance events are hard to detect, yet can significantly affect distributions of genetic variation. Not only do they have the potential to distort cline shape (Szymura and Barton (1991), Fig. 1C, D), but they also require that we use new approaches to infer dispersal rates, estimate selection, and quantify the strength of barriers to gene flow.

In this study, we infer how selection and dispersal have shaped a hybrid zone between two varieties of the snapdragon *Antirrhinum majus* subspecies *majus. A.m. majus* has been a model for understanding trait variation dating back to crossing experiments by Mendel and Darwin (Hudson et al., 2008).

The natural range of *A.m. majus* is mainly centreed in France and Spain, though its popularity in horticulture has seen populations established throughout Europe and across the globe (Hudson et al., 2008). We focus on a natural hybrid zone between two varieties, *A.m.m* var. *pseudomajus* and *A.m.m* var. *striatum* (Rothmaler, 1956). They have largely distinct geographic ranges, but occupy a similar range of local habitats and are pollinated by the same bee species (Whibley *et al*., Tavares *et al*., *2018*). The only known trait that distinguishes them is their different flower colour patterns, which are thought to be alternate solutions to attracting bees and signposting bee entry into the flower (Whibley *et al*., Tavares et al., 2018): *A.m.m* var. *pseudomajus* has magenta flowers coloured by anthocyanin pigment with a small patch of yellow aurone pigment below the bee entry point (Fig. 2A), whilst *A.m.m* var. *striatum* has primarily yellow flowers due to widespread presence of aurone pigment, with restricted veins of magenta anthocyanin pigment above the bee entry point (Fig. 2A, Bradley et al. (2017)). This difference in pattern is determined by a handful of loci that control the production and distribution of anthocyanin and aurone pigment in floral tissue (Table 1). Three loci—*ROSEA* (*ROS*) (Tavares et al., 2018), *ELUTA* (*EL*) (Tavares et al., 2018) and *SULFUREA* (*SULF*) (Bradley et al., 2017)—have major effects, while three others— *RUBIA* (*RUB*) (Field et al., 2025), *CREMOSA* (CRE) (Richardson et al., 2025), and *FLAVIA* (*FLA*) (Bradley et al., 2025) — have more subtle effects on colour pattern and/or intensity.

**Table 1:**
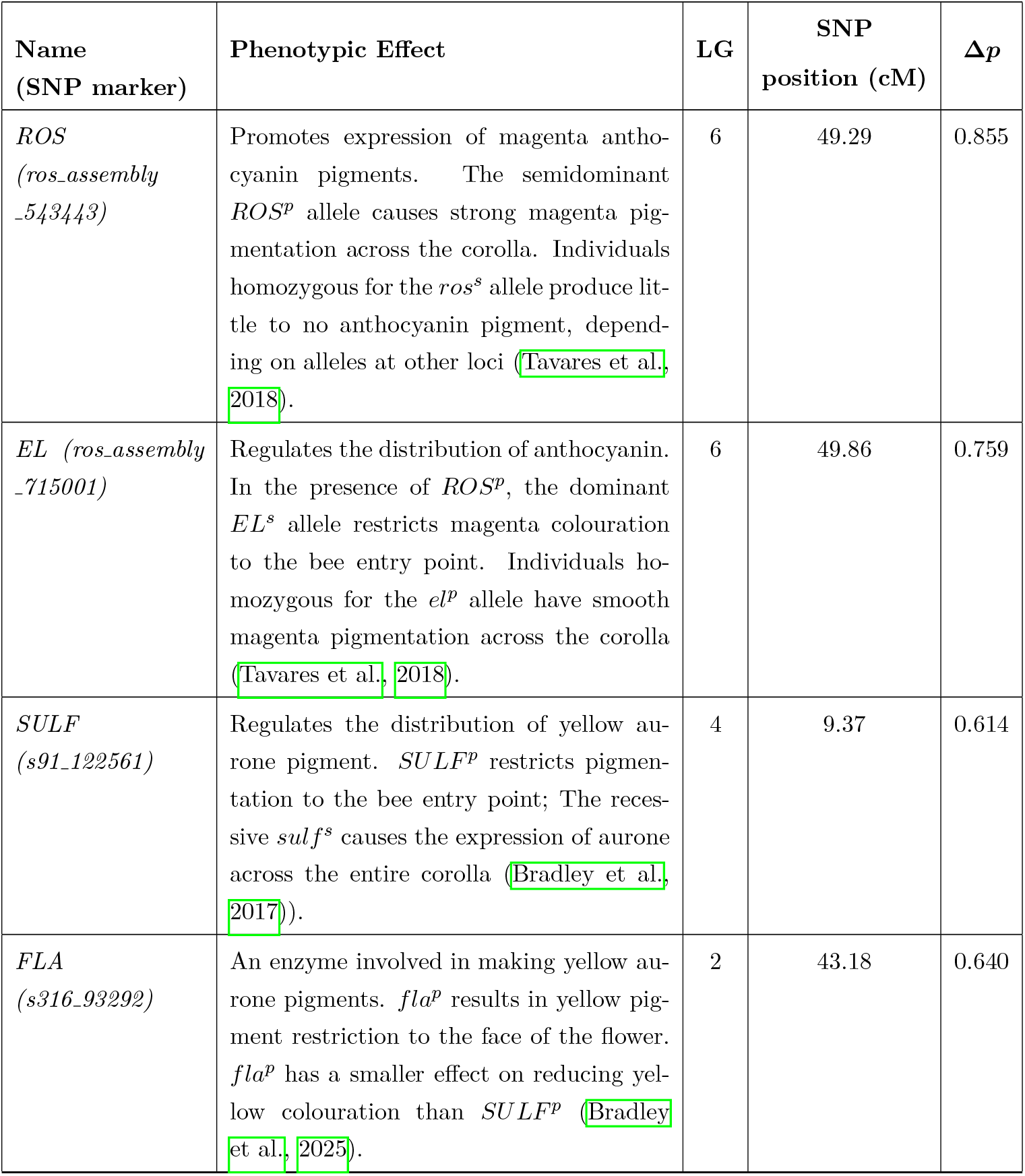
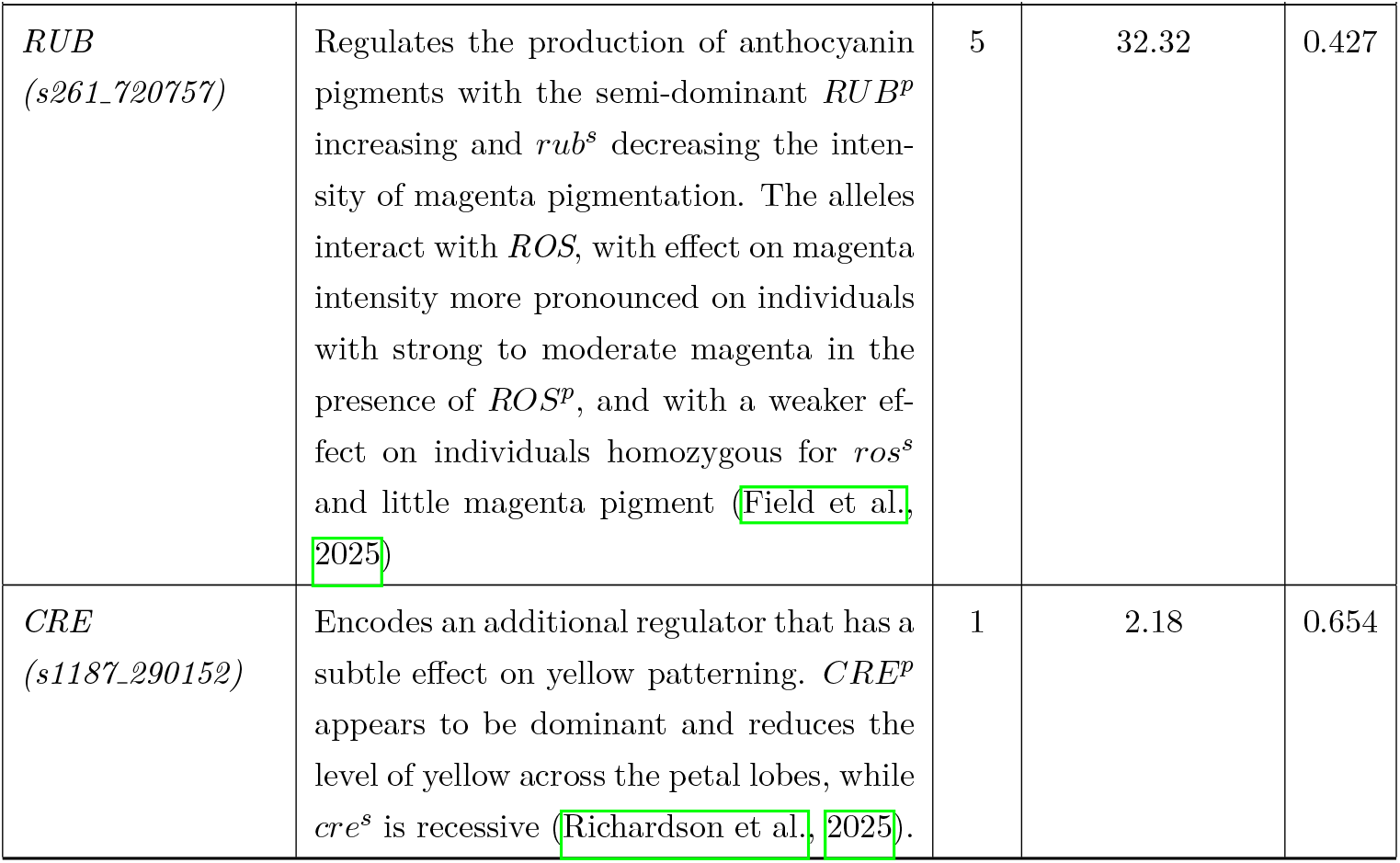
Information about the colour loci and corresponding SNP markers studied, with basic descriptions of the effect of each locus. The superscript letters indicate which subspecies the allele is typically associated with: *p* = *A.m.m* var. *pseudomajus, s* = *A.m.m* var. *striatum*. LG is the linkage group where the locus is found. SNP position (cM) is the position of the genotyped SNP in centimorgans. Δ*p* is the allele frequency difference between the left (yellow) and right (magenta) flanks, defined as *z <* 13 km and *z >* 16 km on the 1-D transect respectively, where *z* is the distance along the transect (see below).

| Name<br>(SNP marker) | Phenotypic Effect | LG | SNP<br>position (cM) | $\Delta p$ |
| --- | --- | --- | --- | --- |
| *ROS*<br>(*ros_assembly*<br>_543443) | Promotes expression of magenta anthocyanin pigments. The semidominant $ROS^p$ allele causes strong magenta pigmentation across the corolla. Individuals homozygous for the $ros^s$ allele produce little to no anthocyanin pigment, depending on alleles at other loci (Tavares et al., 2018). | 6 | 49.29 | 0.855 |
| *EL* (*ros_assembly*<br>_715001) | Regulates the distribution of anthocyanin. In the presence of $ROS^p$, the dominant $EL^s$ allele restricts magenta colouration to the bee entry point. Individuals homozygous for the $el^p$ allele have smooth magenta pigmentation across the corolla (Tavares et al., 2018). | 6 | 49.86 | 0.759 |
| *SULF*<br>(*s91_122561*) | Regulates the distribution of yellow aurone pigment. $SULF^p$ restricts pigmentation to the bee entry point; The recessive $sulf^s$ causes the expression of aurone across the entire corolla (Bradley et al., 2017). | 4 | 9.37 | 0.614 |
| *FLA*<br>(*s316_93292*) | An enzyme involved in making yellow aurone pigments. $fla^p$ results in yellow pigment restriction to the face of the flower. $fla^p$ has a smaller effect on reducing yellow colouration than $SULF^p$ (Bradley et al., 2025). | 2 | 43.18 | 0.640 |
| *RUB*<br>(*s261_720757*) | Regulates the production of anthocyanin pigments with the semi-dominant *RUB<sup>p</sup>* increasing and *rub<sup>s</sup>* decreasing the intensity of magenta pigmentation. The alleles interact with *ROS*, with effect on magenta intensity more pronounced on individuals with strong to moderate magenta in the presence of *ROS<sup>p</sup>*, and with a weaker effect on individuals homozygous for *ros<sup>s</sup>* and little magenta pigment (Field et al., 2025). | 5 | 32.32 | 0.427 |
| *CRE*<br>(*s1187_290152*) | Encodes an additional regulator that has a subtle effect on yellow patterning. *CRE<sup>p</sup>* appears to be dominant and reduces the level of yellow across the petal lobes, while *cre<sup>s</sup>* is recessive (Richardson et al., 2025). | 1 | 2.18 | 0.654 |

**Figure 2.**
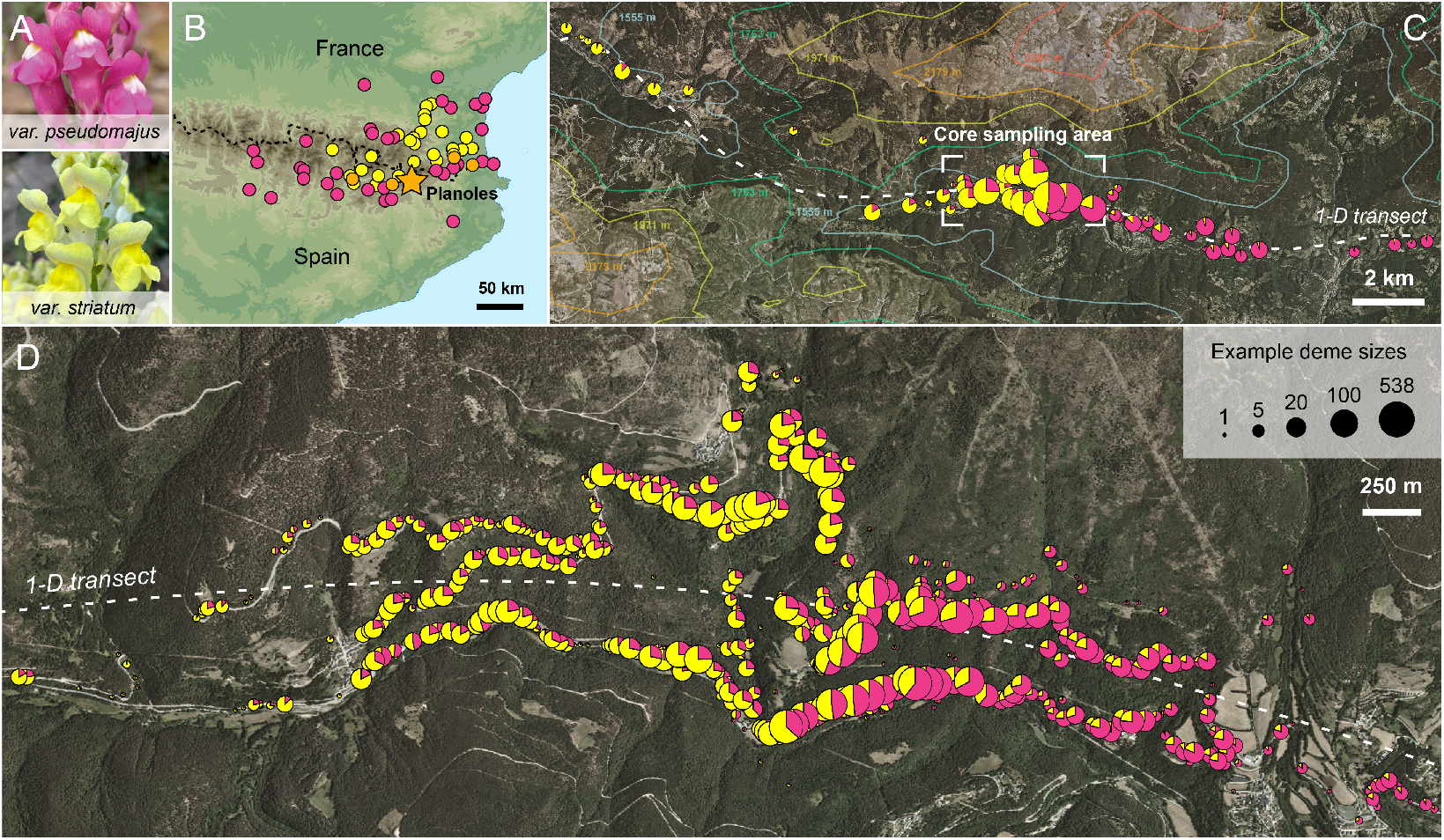
The snapdragon hybrid zone in the Spanish Pyrenees. (A) Typical phenotypes of the varieties. (B) Partial ranges of *A.m.m* var. *pseudomajus* (magenta), *A.m.m* var. *striatum* (yellow) and known hybrid zones (orange) based on sampling in Whibley et al. (2006). The magenta flowered *A.m.m* var. *pseudomajus* is found in northern Spain and south-western France, its range encircling the yellow-flowered *A.m.m* var. *striatum*. The dashed black line marks the French-Spanish border; the star indicates our study area. (C) The hybrid zone in the Val de Ribes. For visualisation, the 22,494 individuals have been clustered into 500m demes; pie charts show the mean hybrid index within each deme, defined as the proportion of *A.m.m* var. *pseudomajus* alleles at five unlinked colour loci (excluding *EL* which is tightly linked to *ROS*). The dashed white line shows the one-dimensional transect line and the contour lines show altitude. (D) The core sampling area. Individuals are clustered into 999 25m demes, which is the scale used for all analyses. In C and D, pie charts are scaled according to the log of the number of plants that they contain.

During the last ice age, *A.m. majus* was restricted to areas of lower elevation, but subsequently expanded into the Pyrenees as the climate warmed. As a result, *A.m.m* var. *pseudomajus* and *A.m.m* var. *striatum* have come into contact, forming narrow hybrid zones in multiple valleys separated by mountains that rise above the altitude tolerance of *A.m. majus* (Fig. 2B). Segregation and recombination of colour alleles in hybrids has generated a wide range of phenotypes. These include individuals that are phenotypically intermediate between *A.m.m* var. *pseudomajus* and *A.m.m* var. *striatum*, as well as unusual phenotypes that fall outside the range seen in the genus Antirrhinum (Tastard et al., 2008). Selection by pollinators is thought to maintain hybrid zones in two ways. First, experiments have shown that some hybrid colour patterns are less likely to be visited because they do not fit the visual preferences of bee pollinators (Tastard et al., 2008). *A.m.m* var. *pseudomajus* and *A.m.m* var. *striatum* colour patterns are also thought to be subject to positive frequency-dependent selection and/or selection against hybrids, each having higher fitness on the side of the hybrid zone where they are more common (Tastard et al., 2014).

Several studies at one hybrid zone in the Val de Ribes suggested that selection acts on flower colour. First, past work has shown that the transition from *A.m.m* var. *pseudomajus* and *A.m.m* var. *striatum* is abrupt, and is characterised by a steep cline in flower colour only a few km wide (Whibley et al., 2006; Tavares et al., 2018; Bradley et al., 2017). Studies of genome-wide sequence variation, including preliminary cline analyses, have revealed strong allele frequency differentiation and steep clines around colour loci (Whibley et al. (2006); Field et al. (2025)). In contrast, the remaining genomic background shows low differentiation, potentially due to the long-term effects of dispersal and recombination between the two subspecies (Whibley et al. (2006); Tavares et al. (2018); Ringbauer et al. (2018); Field et al. (2025); Pal et al. (2025)). Despite this past work, we still lack a robust, quantitative understanding of how selection and dispersal shape this hybrid zone.

In this study, we conduct a genetic analysis of the hybrid zone in the Val de Ribes, based on 22,494 individuals sampled and genotyped over 10 years (Field et al. in prep.). Our main aim was to interpret cline shapes and LD among SNP markers that are tightly linked to 6 known colour loci in order to quantitatively understand how selection and dispersal shape the zone. Our analysis is based on one SNP marker per colour locus; we refer to these by the name of the nearby causal locus (*ROS, SULF*, …), but emphasise that these are only markers for causal loci that may have a complex structure. A novel aspect of our approach, made possible by a recently inferred pedigree for this population (Field et al. in prep) and intensive sampling, is that we are able to make use of the full distribution of pollen and seed dispersal, which is highly leptokurtic. We show that long-range dispersal events which are not accounted for in traditional cline theory, substantially distort the cline shape. By incorporating the empirical dispersal into a multilocus simulation, we explain the causes of stepped clines and observed LD and decompose the total effective selection into direct and indirect components. Our results expand our understanding of the processes that shape this hybrid zone, and highlight new approaches to infer dispersal, selection and barriers to gene flow when the assumptions of ‘traditional’ cline analysis do not match reality.

## 2 Results

### 2.1 Sampling and description of datasets

Our study uses data that was collected as part of a long-term collaborative initiative to survey the hybrid zone started by researchers from the John Innes Centre (Norwich, UK) and the Institute of Science and Technology Austria. We analysed the data collected each year from 2009 to 2019 during the peak of the flowering season (late May to early August), attempting to sample every flowering individual within our core study area (See Methods; Fig. 2C and D). Although the area is large, snapdragons are restricted mainly to south-facing rocky cliffs and disturbed areas that are accessible from two parallel roads. Repeated surveys of adjacent forested slopes, which are heavily shaded and dominated by conifer species of *Pinus* or *Abies*, have found very few individuals, suggesting that we survey most of the suitable habitats in the area. Other suitable habitat includes disturbed deforested areas further from the roads, small side roads, railway embankments, and hiking trails. In some years, we also sampled beyond the core area to improve sampling in the left and right tails of the cline (Fig. 2C). After accounting for individuals that were sampled multiple times (identified by genotype) because they lived more than one year, our dataset includes 22,494 unique individuals used in subsequent analysis.

Our analysis is based primarily on three types of data. First, we have the sampling year and spatial position of each plant. Second, each plant has been genotyped at ∼ 100 SNP markers. Six of the SNPs (used in the subsequent cline analysis) are tightly associated with causal alleles at six of the known colour loci. These include three loci that influence the production and spread of magenta anthocyanin pigment (*ROS, EL* and *RUB*) and three that influence the production of yellow aurone pigmentation (*SULF, FLA, CRE*). Except for *ROS* and *EL*, which are located about 0.5 cM apart on LG6, the loci are on different chromosomes (Table 1). The remaining SNPs were chosen based on their power to identify parent-offspring trios across our temporal dataset. Because the spatial locations of the parents and offspring are known, inferred trios provide us with direct estimates of dispersal in our study area. Third, we have qualitative colour scores for each individual, describing the level of magenta and yellow colouration of one sampled flower. Because these scores are subjective (flowers were scored by different observers over the last decade), we limit our analysis of these scores to a qualitative comparison with the SNP clines. We do not attempt to distinguish alternate forms of selection or analyze phenotypes in detail: we most likely do not have substantial power to distinguish subtly different models despite our very large sample.

### 2.2 Clines at flower colour loci have similar positions but different shapes

We first fit clines to each of the 6 colour loci separately (see Methods, Table 1). To facilitate cline fitting, we first clustered plants into 999 demes and calculated their position along a one-dimensional transect (z) (see Methods). A 25m diameter was chosen because it yielded demes with enough individuals to estimate local allele frequencies and LD (mean of 23 individuals per deme; 337 demes with at least 10 individuals, Table S1), and minimize departure from Hardy-Weinberg equilibrium (Fig. S3). We then used the Metropolis-Hastings algorithm to fit three cline models to the allele frequency data, while describing variation around the cline using *F*_*ST*_ (see Methods). These included (i) a simple sigmoid model, (ii) a sigmoid model with polymorphism in the tails of the cline, and (iii) a stepped model, characterized by a central sigmoid shape flanked by long exponential tails of introgression which may differ to the left and right (Fig. 1B). Analysis of one locus (*ROS*) found no changes in cline position or width over time (Fig. S4, Table S3), so we combined individuals across all years. We compared likelihoods to identify the best-fitting model for each locus, and obtained maximum-likelihood estimates (MLE) for each cline parameter (Table 2).

**Table 2:**
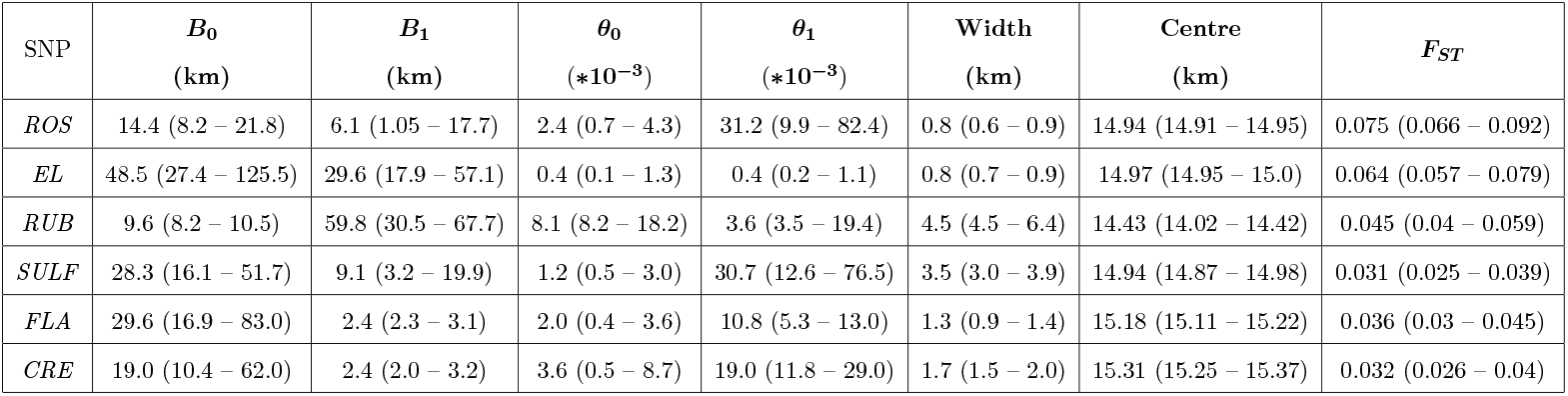
Maximum likelihood estimates and support limits of the 7 cline parameters for the stepped cline model for each flower colour SNP marker. The cline width is the inverse of the maximum gradient in allele frequency, the cline centre denotes the position on the transect at which both alleles are equally frequent and *F*_*ST*_ captures variation in allele frequency around the cline. *B*_0_ and *B*_1_ can be interpreted as the strength of the barrier to gene flow to the left (*striatum* side) and right (*pseudomajus* side) of the cline centre, and *θ*_0_ and *θ*_1_ are the rate of decay of the exponential tail to the left and right sides, respectively.

All six of the colour loci show sharp clines which match the transition in flower colour scores (Fig. 3A, B). The stepped cline model fits the data better than both sigmoid models for all 6 loci (Table S2, Fig. 3B). We examined the MLE (see Table 2) and the marginal likelihood (equivalent to the posterior distribution) to see how parameters differed among loci. The cline centres were highly similar among loci (Fig. 3C, Table 2), ranging from 14.43 km to 15.3 km (mean c = 14.96 km), and differing from one another by an average of 0.34 km (maximum of 0.9 km). Cline widths, on the other hand, differed markedly among loci (Fig. 3C), ranging from 0.76 km (*ROS*) up to 4.5 km (*RUB*). We also found striking differences in the degree of polymorphism and introgression in the tails of the cline. While most SNP markers fix alternate alleles on either side, some loci showed appreciable polymorphism in the yellow flank (i.e., *RUB* and *SULF*, where the *pseudomajus* allele has a frequency around 0.2). Additionally, *FLA* and *CRE* showed strong asymmetries in the cline shape with rates of decay of tails of introgression that are much higher on one side (e.g., *θ*_1_ *> θ*_0_).

**Figure 3.**
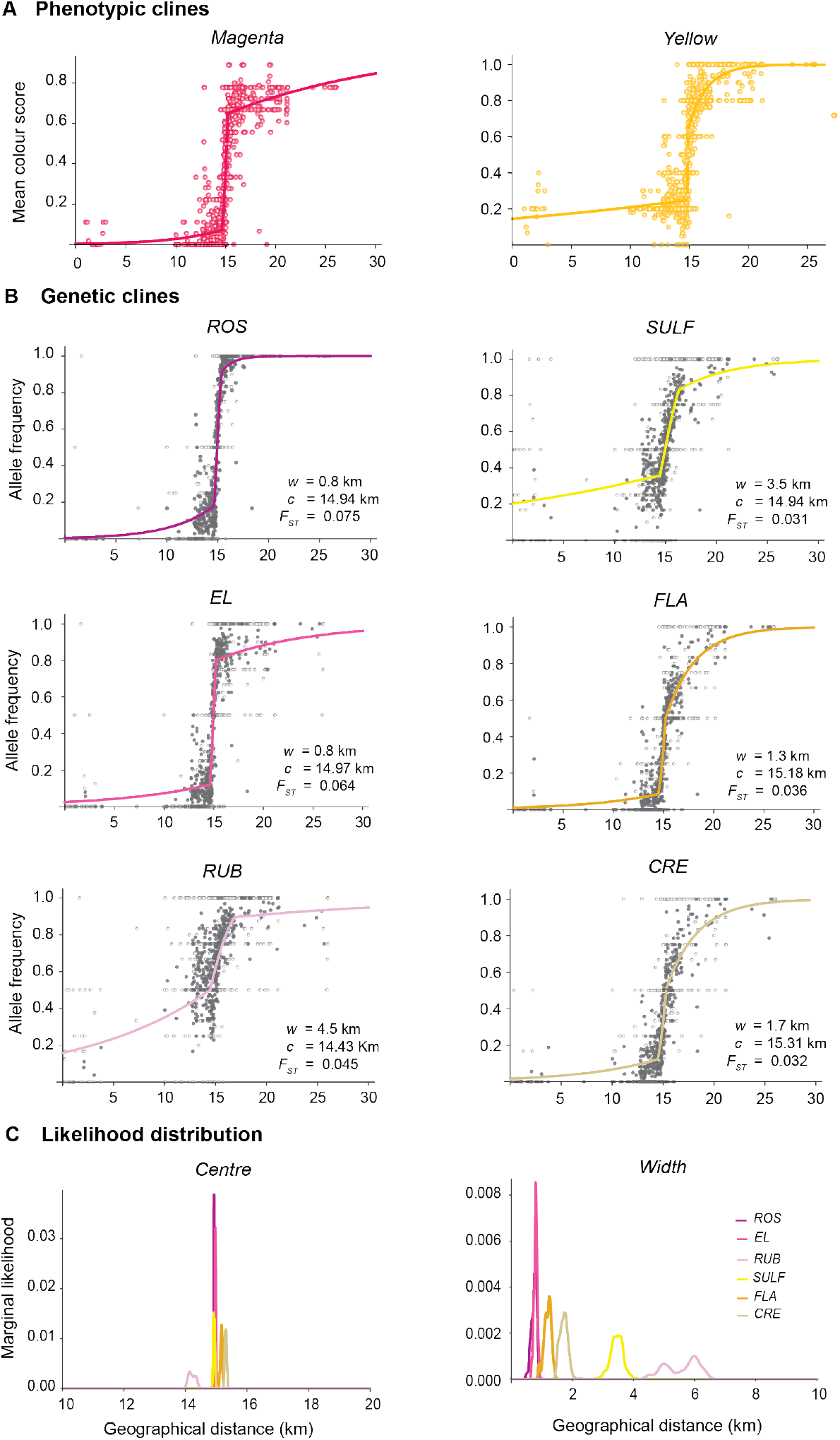
Stepped clines in snapdragons. (A) Phenotypic clines for the mean magenta and yellow colour scores across 999 demes in the hybrid zone. The open circles denote the mean colour scores in each deme and the coloured curve represents the best fit cline from Metropolis-Hastings algorithm. (B) Best fit stepped cline shape and parameters, including the cline width (*w*), cline centre (*c*), and *F*_*ST*_, for the 6 flower colour loci. Gray and light gray circles denote demes with greater and less than 5 individuals respectively. The curves show the asymmetric stepped cline shapes fitted to the observed allele frequencies for all loci. (C) The marginal likelihood of cline centre and width for all 6 loci. Clines roughly coincide, with all cline centres positioned within 1 km. Cline widths vary, with *ROS* narrowest and *RUB* widest. Here, the 1D marginal likelihood distribution is obtained from the last 2500 runs of the Metropolis algorithm.

Taken together, and in the context of the low genome-wide differentiation across this hybrid zone (Tavares et al. 2018), our results are consistent with the hypothesis that the colour loci are targets of pollinator-mediated selection. The varying widths indicate that some loci experience stronger selection than others, likely owing to their individual effects on colouration and/or variation in their degree of linkage to causal mutations. However, the similar positions of the cline centres are expected, given that the difference in colour pattern between *A.m.m* var. *pseudomajus* and *A.m.m* var. *striatum* is determined by interactions between these loci. This coincidence of clines may in addition be due to the presence of associations amongst the six loci, which we examine next.

### 2.3 Associations between flower colour markers extend beyond the cline centre

Studies of hybrid zones often find strong associations between alleles at unlinked loci (i.e., linkage disequilibrium, LD) in areas where clines coincide (e.g., *Bombina*, Szymura and Barton (1986, 1991); *Heliconius*, Mallet et al. (1990); *Vandiemenella*, Kawakami et al. (2009), *Mus musculus*, Macholán et al. (2007)). The LD is usually attributed to the influx of distinct multilocus genotypes as parental individuals disperse into the zone from both sides (Barton, 1979b; Barton and Shpak, 1990). Because LD among unlinked loci is halved each generation by recombination, it should decay rapidly with increasing distance from the cline centre under short-range dispersal.

To test for associations in the hybrid zone, we calculated standardised LD within each deme for the five unlinked loci (excluding *EL*, noting that choosing *ROS* or *EL* makes little difference due to their tight association). We used an LD estimator,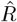, based on the variance of hybrid index (HI). HI is defined as the number of *pseudomajus* alleles carried by each individual, such that with 5 loci, HI ranges from 0 to 10. The mean HI in each deme, calculated from all 5 loci, is plotted along the transect in Fig. 4A; it shows a clear stepped pattern and has a shape and position qualitatively similar to the single locus clines. The variance of the hybrid index, *V*_*HI*_ = 2∑ _*i*_ *p*_*i*_*q*_*i*_ + 2∑ _*i*_ *p*_*i*_*q*_*i*_*F*_*IS,i*_ + 2∑ _*i≠j*_ *D*_*ij*_, separates into components due to heterozygosity (*V*_*P Q*_), heterozygote deficit (*V*_*F*_), and linkage disequilibrium respectively (modified from Barton and Gale (1993)). Both LD and heterozygote deficit inflate the variance in the hybrid index, meaning that the excess variance can be used to estimate the standardised LD or correlations relative to linkage equilibrium given as 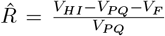. This was calculated separately for each pair of loci (maximum likelihood gives similar results); 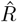 ranges from -1 to 1. We also calculated heterozygote deficit, *F*_*IS*_, for each locus within each deme. To reduce noise among demes, we calculated mean 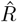 and *F*_*IS*_ in 12 non-overlapping bins along the transect, and 3 coarser bins to examine LD inside and outside the cline centre.

**Figure 4.**
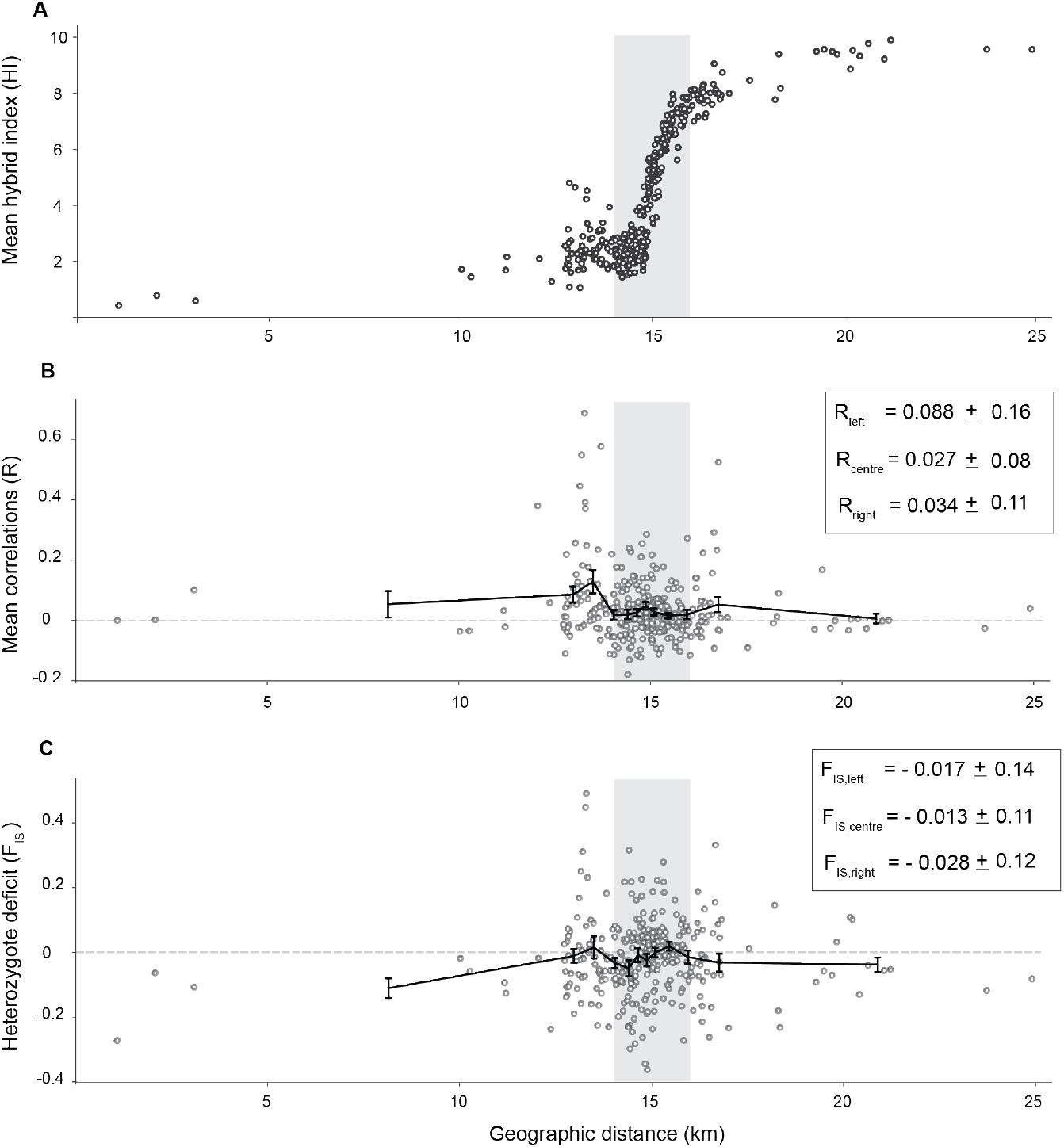
Little heterozygote deficit and positive correlations across the transect. (A) Mean hybrid index from 5 unlinked loci shown across geographical distance for 337 demes with at least 10 individuals. (B) Mean correlations 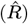 for all 10 pairs of unlinked loci across geographic distance and (C) Mean heterozygote deficit for the 5 loci (*F*_*IS*_). Each point is the estimate for 1 deme, while the black line represents the mean and standard deviation for the 12 bins across the transect. The legends in B and C show the mean *F*_*IS*_ and 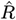 in the centre (denoted by grey bar), magenta flank (right of the grey bar) and yellow flank (left of the grey bar).

Combining individuals from the centre and flanks, we observed significantly positive correlations between all but one pair of loci (permutation test, see Methods). *ROS* had the strongest associations with *FLA* (0.089) and weakest with *RUB* (0.029), while *RUB* and *SULF* showed weak LD (−0.002) (see Table S7, Fig. S6). However, the mean correlations 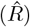 across all pairs of loci were variable and non-zero across the 12 bins along the transect, ranging from 0.006 to 0.12 (Table S6). 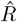 was significantly positive but modest in the cline centre, averaging 0.027 across the central 2km region (Fig. 4B). Additionally, we found no evidence for a marked deficit of heterozygotes at the cline centre, as most bins had a mean *F*_*IS*_ near 0, after averaging over the 5 loci (Table S6, Fig. 4C). However, *ROS* showed a deficit of heterozygotes (*F*_*IS*_ = 0.034) possibly due to strong selection or assortative mating (Table S5, Fig. S5). Overall, the low levels of LD and *F*_*IS*_ show that there is only a weak genome-wide barrier, consistent with Ringbauer et al. (2018), which found no evidence of a barrier to gene flow from non-clinal SNPs.

Traditional cline analysis allows the inference of dispersal and selection from associations between loci (LD) and cline widths (Barton and Gale, 1993). Mean correlations in the centre of the hybrid zone can be used to estimate dispersal as 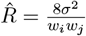, where *w* is the cline width of locus *i* and *σ*is the root mean square of parent-offspring dispersal distance along the transect. The mean correlations between all pairs of unlinked loci in the centre, 0.027, implies *σ*= 98m. This estimate of dispersal can in turn be used to infer the strength of selection from 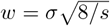, assuming selection against heterozygotes (Bazykin, 1969). Using MLE for cline width (Table 2), estimated selection coefficients ranged from 0.12 in *ROS* to 0.004 in *RUB* (Table 3). This is the effective selection that is required to maintain the cline of the observed width. However, these inferences are valid only when the assumptions of basic cline theory are met.

**Table 3:**
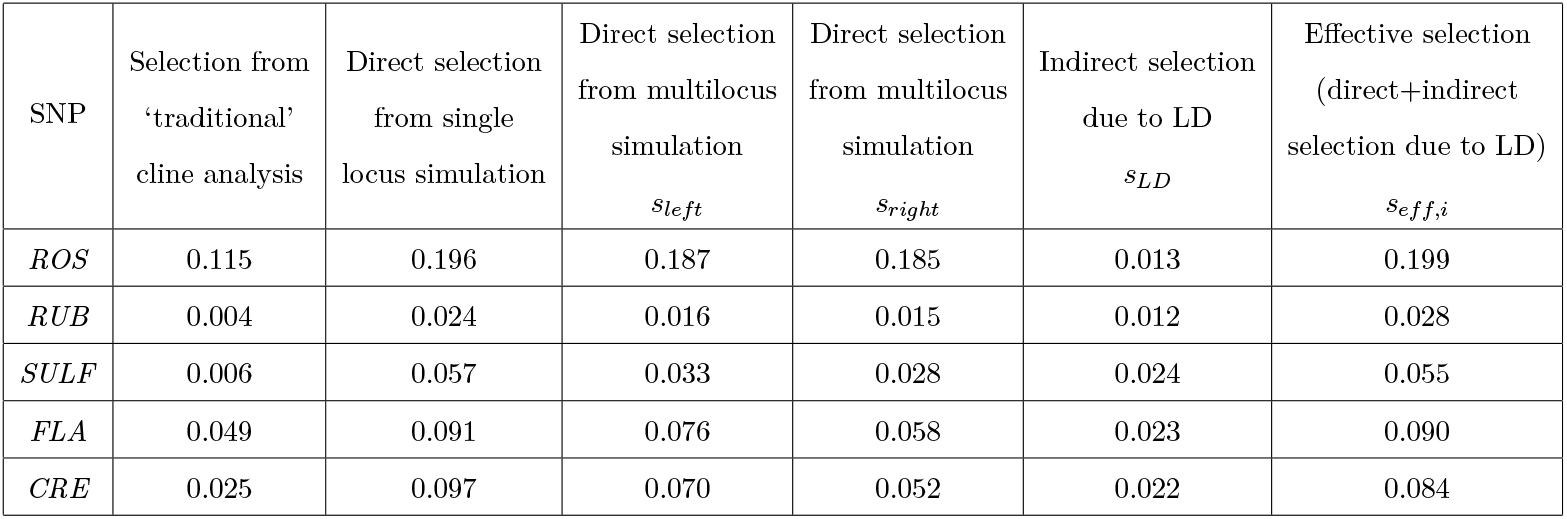
Inferred selection coefficients from cline fits, single locus cline simulations and multilocus cline simulations with asymmetric selection for the 5 unlinked loci. *s*_*left*_ and *s*_*right*_ are the selection coefficients for the two homozygotes respectively. *s*_*LD*_ is the indirect selection (at the cline centre) at each locus due to its associations (LD) with other selected loci. The last column is the effective selection (*s*_*eff,i*_) at each locus at the cline centre due to both its direct effect on fitness and the indirect effect of LD with other selected loci.

| SNP | Selection from<br>'traditional'<br>cline analysis | Direct selection<br>from single<br>locus simulation | Direct selection<br>from multilocus<br>simulation<br>$s_{left}$ | Direct selection<br>from multilocus<br>simulation<br>$s_{right}$ | Indirect selection<br>due to LD<br>$s_{LD}$ | Effective selection<br>(direct+indirect<br>selection due to LD)<br>$s_{eff,i}$ |
| --- | --- | --- | --- | --- | --- | --- |
| *ROS* | 0.115 | 0.196 | 0.187 | 0.185 | 0.013 | 0.199 |
| *RUB* | 0.004 | 0.024 | 0.016 | 0.015 | 0.012 | 0.028 |
| *SULF* | 0.006 | 0.057 | 0.033 | 0.028 | 0.024 | 0.055 |
| *FLA* | 0.049 | 0.091 | 0.076 | 0.058 | 0.023 | 0.090 |
| *CRE* | 0.025 | 0.097 | 0.070 | 0.052 | 0.022 | 0.084 |

Our results highlight two major discrepancies from traditional cline theory and the necessity to adopt a different method, as detailed below. Firstly, the standardised LD in the centre of the hybrid zone is weak (∼0.027). In our system, relatively few loci are under selection (Tavares et al. (2018)), and we show below that the weak LD among them is insufficient to cause a detectable step, as opposed to scenarios where strong LD reflects a genetic barrier to gene flow (see Fig. S13). Stepped clines can also reflect a geographical barrier to dispersal (Barton and Hewitt, 1985; Westram et al., 2022); however, this seems unlikely, as we see no indication of a strong geographical barrier in this hybrid zone. It is also possible that the stepped shape is caused by introgression of a marker SNP as it recombines away from a tightly linked causal locus (Barton, 1979b); we return to this issue in the Discussion.

A second unexpected finding is the presence of positive LD across the entire transect (Fig. 4B); in fact, the correlation between alleles is higher in the flanks than in the centre (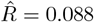 in the yellow flank and 0.034 in the magenta flank, compared to 0.027 in the centre). The simplest explanation is that this is generated by long-distance dispersal from one side of the hybrid zone to the other (Szymura and Barton, 1986, 1991). Long-distance dispersal directly brings in sets of foreign alleles generating introgression and causing local increases in LD away from the cline centre. In such scenarios *σ*may not suffice to describe dispersal: the diffusion approximation fails, and inferences from traditional cline analysis may be inaccurate.

### 2.4 Snapdragons show extreme long-distance dispersal

To understand how dispersal influences the cline shape and LD, we inferred the full pollen and seed dispersal distributions from our long-term dataset. Dispersal up to ∼1km (hereafter referred to as short-range dispersal) was obtained using the parent-offspring distances from the 2342 parent-offspring trios inferred in Field et al. in prep. Although this dispersal is leptokurtic, here we refer to the full dispersal distribution-spanning the range of the hybrid zone – which is even more strongly leptokurtic due to the presence of rare long-distance dispersal events far into the tails of the cline.

Beyond the range of the pedigree, we identified long-range migrants in the flanks as having multilocus genotypes which are unlikely to be produced locally by random mating, given the local allele frequencies (SM Text S3). Using this approach, we found that 3.7% and 1.1% of individuals in the yellow and magenta flanks respectively were unlikely to be the result of mating between two local parents (*p <* 2 * 10^*−*4^; Table S8). We infer that these ‘improbable individuals’ arose primarily through longrange seed dispersal. This is because pollen dispersal would yield F1-like individuals in the flanks, yet these are relatively rare (Table S8); moreover, from the pedigree trios (Field et al. in prep.), seed dispersal is more frequent in the extreme tails (beyond the range shown in Fig. 5A). Therefore, we extrapolate over longer ranges from the shape of the short-range pollen dispersal kernel, which closely matches a log-Gaussian distribution (Fig. 5A).

**Figure 5.**
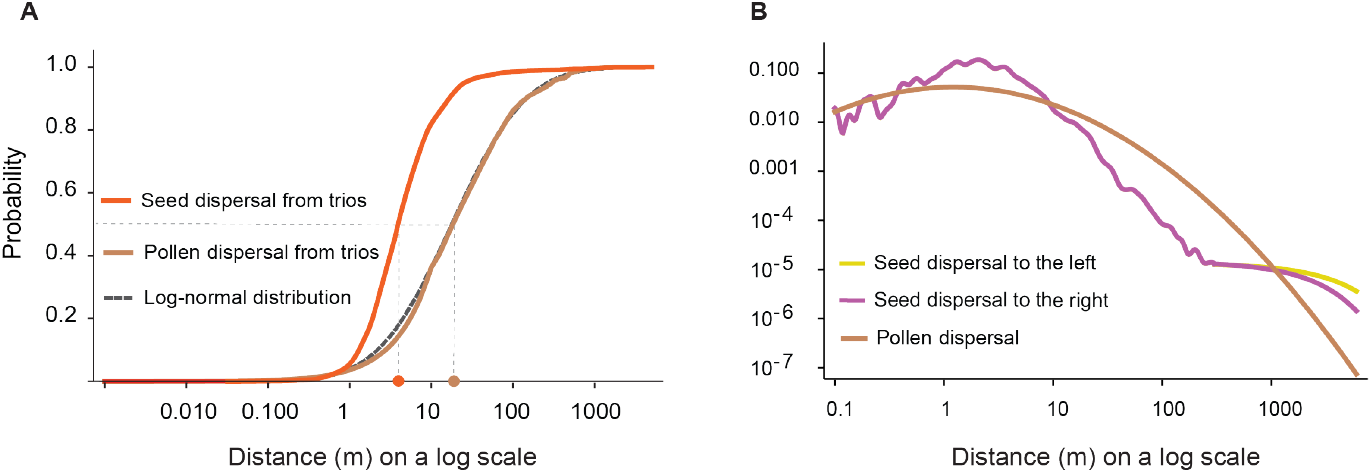
Dispersal is leptokurtic is snapdragons. (A) Cumulative distribution (CDF) of seed and pollen dispersal estimated from the pedigree trios. The brown curve denotes the CDF of the pollen dispersal (i.e. the distance between the parents). The black dashed line denotes the fitted log-Gaussian distribution. The orange curve denotes the CDF of seed dispersal, estimated from the trios. For each curve, the grey dashed lines denote the median distance, where the probability is 50%. (B) The full seed and pollen dispersal distributions, with the probability of movement of a given distance shown in the Y axis. The brown curve denotes the full pollen dispersal kernel fitted to a log-Gaussian. The magenta and yellow curves denote the seed dispersal kernel estimated by splicing the PDF estimated from the trios (as shown in A) with that estimated from individuals in the magenta and yellow flanks respectively at 300m.

In contrast, we inferred long-range seed dispersal separately for individuals in either direction from the improbable parental genotypes that we see in the flanks. We assumed an exponential function which was spliced onto the short-range distribution at 300m, to give a full seed dispersal kernel, fitting the rates of decay by maximum likelihood (see Methods, Fig S8).

Together, these approaches showed that dispersal is highly leptokurtic. Half of fathers were within 18m of the mother, 75% within 55m, and 93% within 200m (Fig. 5A). Seed dispersal was more leptokurtic, with ∼60% of dispersal occurring within 5m and ∼80% within 10 m (Fig. 5A), but with very long tails in both directions (Fig. 5B); the scale of decay in the tails was inferred to be *λ*_*left*_= 4.5km to the left flank and *λ*_*right*_= 2.6 km to the right. Long-range seed dispersal was higher from magenta to yellow, consistent with the high rate of introgression seen from the hybrid index score and greater LD observed in the yellow flank (Fig. 4A,B). These long-range seed dispersers can directly bring in foreign genotypes, thereby generating positive associations between alleles at different causal loci away from the cline centre.

### 2.5 Using simulations with empirical dispersal to explain cline shape and LD

We observe associations (LD) even between unlinked loci, and see strongly leptokurtic dispersal. In order to find the relative contribution of these to distorting clines away from a simple sigmoid, and to estimate the strength of selection that best explains our data, we simulate the five unlinked loci, using the observed dispersal kernel (We exclude *EL*, but note that due to tight linkage between *ROS* and *EL*, simulating *ROS* includes the effect of both these loci; Tavares et al. (2018)), using the observed dispersal kernel. There are a plethora of possible models for selection: with 5 loci, we have 243 diploid genotypes, which can each be assigned a fitness that depends on the frequency of all the genotypes. We are skeptical that we have much power to estimate much more than the net selection on each locus, and so choose as simple a model as possible, at least for this first analysis. We assume random mating, so that diploid genotype frequencies can be constructed from the frequencies of the 2^5^=32 haplotypes. We assume an infinite population, using a deterministic simulation that neglects drift. We also assume that selection is multiplicative across loci, and acts against heterozygotes; it may be asymmetric, so that contribution to fitness at a locus is 1 + *s*_*left*_: 1: 1 + *s*_*right*_. Though we simulate haplotype frequencies, we find the best fit to the allele frequency clines. Because the clines are strongly stepped, we use unevenly spaced demes, so that we can follow steep gradients in the centre, and shallower tails out at the edges. The migration matrix is constructed from the CDF of the observed dispersal, to give the fraction in each ‘deme’ that derives from each of the other demes in every generation. This scheme should be seen as a surrogate for a range of more detailed models; for example, selection against heterozygotes is equivalent to positive frequency-dependent selection, or to epistasis (Bazykin, 1973; Barton, 1979a). Essentially, the selection coefficients give the rate of change of allele frequency on either side, and represent marginal values that absorb the effects of dominance, epistasis and frequency-dependence. Models will differ in the pattern of linkage disequilibrium, but because loci are unlinked, we have littler power to detect such affects, and do not attempt to fit linkage disequilibrium.

Clines that are maintained by selection against hybrids, or by positive frequency-dependence, are not tied to any particular location, and so will tend to move. Such movements are likely to be countered by small density gradients; assuming diffusion, a wave speed *v* can be balanced by a gradient in log density *v/σ*(Barton, 1979a). Therefore, we shift the simulation such that it is always centred in the same place, and find its most likely location with respect to the observed allele frequencies. When we simulate a single locus, asymmetric selection is redundant, since it just shifts the cline without altering its shape (Barton, 1979a). However, when we simulate multiple loci, they can be held together by LD, so that differences in asymmetry between loci cause differences in position, as well as differences in the rate of decay of tails of introgression on either side. Simulated allele frequencies were fit to the observed allele frequencies after 500 generations, to infer MLE of the strength of selection, cline centre, and *F*_*ST*_. We first validated the simulation by considering Gaussian dispersal with *σ*=200m and *s*= 0.1 and following the allele frequencies at a single locus. As expected, we get a sigmoid cline with width 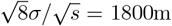 (see Fig. S8).

To understand how long-range dispersal affects cline shape in the absence of LD, we first simulated each locus separately. Simulated clines with only the short-range dispersal (i.e the dispersal inferred in Field et. al (in prep.) from the pedigree trios, up to 300m) failed to generate a strong step (Fig. S9). However, inclusion of long range dispersal generated stepped clines at all loci (log *L* = −9385.3 with 15 df, Fig. S10, Fig. 6A). The inferred selection coefficients for the 5 unlinked loci ranged from 0.196 to 0.024 (Table 3). The simulated clines matched the observed step and patterns of introgression in the tails for *ROS* but failed to explain the complete cline shape at other loci (Fig. S10, Fig. 6A). We next simulated clines at 5 unlinked loci, allowing asymmetric selection, which causes differences in the cline position and rate of introgression. Asymmetric multilocus selection, including long-range dispersal, best explained the observed stepped clines at all loci (log *L* = −9257.8 with 12 df, *F*_*ST*_ = 0.067, Fig. 6A). The inferred selection estimates (see Table 3) were roughly symmetric for *ROS, SU LF* and *RUB*, and differed only by ∼0.02 between the two homozygotes for *FLA* and *CRE*. Similarly, cline centres from cline fitting (*c* = 14.96km) matched closely with that from the simulations (*c* = 14.97 km). However, this simple model does not capture the complete cline shapes, especially at *F LA* and *CRE*.

**Figure 6.**
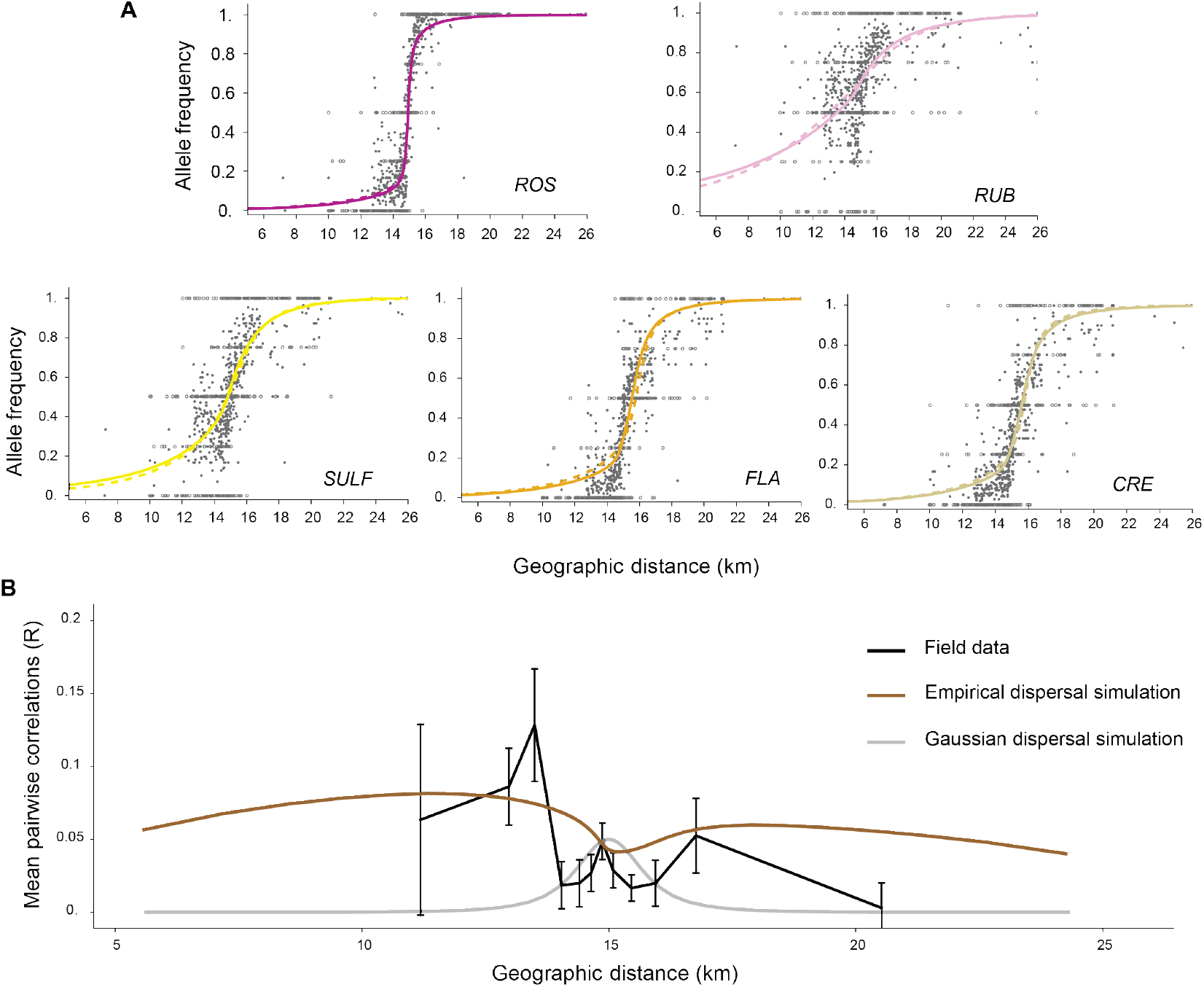
Multilocus cline simulations with the inferred dispersal explain the step and LD. (A) Cline shapes inferred from simulations with asymmetric selection and full dispersal kernel for each locus. The dashed line is the best fit simulated cline from single locus simulations whereas the solid line represents the fit from multilocus simulations with asymmetric selection. The corresponding selection estimates are given in Table 3. The light grey and dark gray circles denote the observed allele frequencies of demes with at least 5 and greater than 5 individuals respectively. (B) Mean correlations from 10 pairs of unlinked loci in the observed data (black) are shown for the 12 bins across the whole transect. The brown and grey curves show the mean correlations from multilocus simulations with the inferred dispersal and Gaussian dispersal (with *σ*=161m) respectively.

We next compared the observed standardized LD with that seen in multilocus simulations. We found that LD tended to be overestimated by the simulation. One cause of this discrepancy might be dependence of LD on the life stage at which it is measured, which in the simulations is immediately after populations mix. However, there is good agreement between the simulated and observed LD patterns (Fig. 6B): mean LD from the simulations was positive across the entire transect, and stronger in the tails of the cline 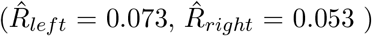 than at the centre 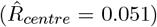. We also found that the strength of the simulated pairwise LD varied as expected, with stronger LD among pairs of loci under stronger selection (Fig. S11). These results confirm that long-range dispersal can generate both widespread LD and stepped clines.

### 2.6 Inferences of the strength of selection and the barrier to gene flow

The multilocus simulations allow us to separate the direct, indirect, and total effective selection acting on each locus. The total effective selection at any locus *i* (*s*_*eff,i*_) is the sum of direct selection experienced by that locus individually (*s*_*i*_), and the indirect selection due to associations with other selected loci (*s*_*LD,i*_), and can be calculated in the cline centre (assuming clines coincide) as 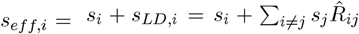, where *s*_*i*_ = (*s*_*left*_ + *s*_*right*_)*/*2 is the average selection coefficient from asymmetric multilocus simulations at locus *i* and 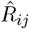 is the correlation between locus *i* and *j*. This varied from 0.199 to 0.028 among the 5 loci at the centre of the hybrid zone (Table 3). The direct estimates of selection (taken as the average of *s*_*left*_ and *s*_*right*_ for asymmetric selection) were lower than from the single locus simulations. This is due to the effect of LD (indirect selection) which increases the total effective selection experienced at a locus, thus lowering the inferred direct selection.

Effective selection at each locus was similar to that from single locus simulations. Multilocus selection has a weak effect on cline shape: LD due to indirect selection only causes a moderate increase in the step (Fig. 6A, Fig. S10). This effect of LD is captured by the indirect selection on each locus, *s*_*LD*_ and its relative contribution to effective selection is *s*_*LD*_*/s*_*eff*_. For each locus, the indirect selection contributed roughly 30% (on average) of the total effective selection. It was weakest in *ROS* (6.7%) and strongest in *RUB* and *SULF* (44%) as expected: the effect of LD is strongest on weakly selected loci.

The multilocus simulation scheme also allows us to gain insight into the strength of the barrier to gene flow acting in this system. In traditional cline analysis, the strength of the barrier to gene flow at the centre of the hybrid zone is estimated from the shape of stepped clines. Because we know that long-distance dispersal and LD contribute to the step, we must first estimate how much of the step is due only to LD. To do this, we conducted an additional multilocus simulation that assumed Gaussian dispersal. We used a dispersal rate of *σ*= 161m estimated from 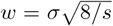, with the cline widths (*w*) estimated from descriptive cline fits and the selection coefficients (*s*) inferred from the multilocus simulation. Although this scenario generated a sharp peak of LD at the cline centre 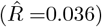 which decayed rapidly with increasing distance (gray curve in Fig. 6B), it generated a negligible step (Fig. S13). Thus, while there must be some barrier due to LD, it is likely negligible and below the limits of detection using theory based on cline shape. This further emphasises the importance of long-distance dispersal in generating the stepped clines.

Although we were unable to detect a barrier at the cline centre via cline fitting, the net barrier to gene flow across the hybrid zone can still be measured by the ratio *B* = Δ*p/p*^*′*^, but it must be measured across the whole region that is affected by selection, which spans ∼10km, and this is difficult, even with our very large sample. Gene flow across this hybrid zone consists of two components: diffusion of genes through the central region, and long-range dispersal, which introduces parental genotypes on either side. The barrier due to diffusive gene flow at an unlinked neutral loci is proportional to the central gradient and is determined by the net selection against hybrids. This can be measured by 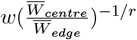, where *w* is the cline width, 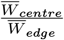 the mean fitness of the hybrid population at the centre relative to the parental type and *r* = 1*/*2 is the recombination rate (Barton 1986). Since mean fitness of the parental types across the cline is 1, this is given by 2 (1 − *s*_*i*_*/*2)^2^ ∼ 0.71. Thus, this component of the barrier reduces gene flow by 30%.

We must also consider the strength of the barrier to gene flow generated by selection against longrange migrants. Here, we obtain a rough estimate by treating the yellow and magenta flanks as discrete demes that exchange migrants at a low rate (*m*). Under this model, the rate of gene flow for an unlinked neutral locus in the presence of selection (*m*_*e*_) relative to that expected without any barriers to gene flow (*m*) is the product of the mean fitnesses of incoming migrants and all successive hybrid classes: *m*_*e*_*/m* = *W*_*m*_ * *W*_*F* 1_ * *W*_*bc*1_ … (Westram et al. *(2022)). In our study, the fitness of each successive migrant can be calculated from our multilocus simulations using the average selection coefficient for each locus. This yields an estimate of m*_*e*_*/m* = 0.5. Thus, the rate at which an incoming neutral migrant allele associated with the opposite phenotype gets established is half the rate at which they enter through long distance dispersal. In other words, the strength of reproductive isolation (*sensu* Westram et al. (2022)) *due to selection acting against long range migrants (1 − me/m*) is 0.5. Thus, both components of the barrier reduce gene flow to a similar degree, but there is a stronger barrier to long-range dispersal, because their offspring suffer stronger selection than do sets of genes flowing through the centre.

## 3 Discussion

Our analysis of the genotypes of 22,494 individuals provides new insights into the processes shaping the flower colour clines in this hybrid zone. We found patterns typical for hybrid zones, including stepped geographic clines at all six colour loci (Fig. 3B), and significant LD at the cline centre (Fig. 4B). However, some of our results were not predicted by traditional cline theory, including positive LD across the entire transect, and stepped clines despite weak LD in the centre. A multilocus simulation allowed us to estimate both direct and indirect selection, and to show that the stepped clines are caused primarily by long-distance dispersal rather than reflecting a strong genetic barrier to gene flow. Here, we elaborate on some of the key conclusions and highlight the general implications of our work for the study of hybrid zones.

Our study shows that these flower color clines are maintained by a balance between dispersal and selection. We also found that the strength of selection obtained from simulations varies among the colour markers, with estimates of direct selection ranging from 1.5% to 19% (Table 3). We think that this variation is mostly due to the different effects that each locus has on flower colour. For example, the tightly linked loci *ROS* and *EL*, which have the largest selection coefficient (19%), are the major determinants of magenta colouration. In contrast, *RUB*, which has the smallest selection coefficient (1.5%), has a relatively modest effect on colour (Field et al., 2025), subtly increasing the intensity of magenta if it is already present (Table 1). Although this interpretation makes intuitive sense, because pollinator preferences are thought to be driven by the differences in flower colour, other factors also contribute to variation in our estimates of selection. First, our SNPs may be incompletely associated with causal mutations that influence the pattern of flower colour (as is usually the case for genetic markers used to study local adaptation). Since their association with the underlying causal loci is almost certainly incomplete, we will inevitably underestimate the strength of direct selection. For example, multiple maker SNPs have been developed around the *SULF* locus. We analysed the one with a broader cline, whilst the SNP analysed by Bradley et al. (2025) shows a narrower cline and a closer association with yellow score, but also has much more missing data (which may be due to a deletion in this region; see Fig. S14 for comparison of cline shapes between these markers). This partly explains the difference in the estimates of selection for SULF between these studies. Thus, we may be underestimating selection at this locus due to our choice of SNP. Second, the causal loci themselves are complex, and may involve multiple causal sites, and structural variation. There are several coding genes around *ROS* and recombinants with *EL* are found at appreciable frequency (Tavares et al., 2018); similarly, at *FLA*, a recombinant haplotype is found between the two parental populations (Bradley et al. (2025), Fig. 4H). We now have whole genome sequences for a smaller sample of ∼1000 individuals from across the cline. In future work, we aim to fit a model using the set of clines surrounding one (or more) selected sites, reflecting the flow of haplotypes past a local genetic barrier.

A unique feature of our study, critical to our main findings, is that we inferred the full dispersal distribution by combining parent-offspring trios (from Field et. al, in prep) with identification of long-distance migrants. This showed that dispersal was highly leptokurtic with primarily short range dispersal, but also a long tail of dispersal that extends beyond 1km (Fig. 5B). Leptokurtic pollen dispersal is commonly observed in bee-pollinated plant species, and has already been documented over a smaller scale in a paternity study in snapdragons from our study area (Ellis et al., 2024). The bees that pollinate snapdragons (*Bombus* spp.) tend to forage locally, but are routinely recorded moving longer distances well beyond 1 km (Walther-Hellwig and Frankl, 2000; Osborne et al., 2008). We found that the seed dispersal distribution was also leptokurtic, with a small fraction of dispersal events exceeding 1 km. Although the vector for long range seed dispersal is not clear, the patchy distribution of the species requires long distance seed dispersal. Moreover, snapdragons are often observed growing on the roofs of buildings and high on the stone walls of churches and castles, also suggesting that long distance seed dispersal is quite common. Leptokurtic dispersal is not unique to snapdragons, yet in most cases, direct estimates of the full dispersal distribution, such as those that come from our pedigree, are not feasible in short-term studies. Nevertheless, it may be possible to estimate both short and long-range dispersal from LD in the centre and in the flanks of the hybrid zone, and then use these to estimate the strength of selection required to explain clines in allele frequency. While this approach would require large samples both within and outside the hybrid zone and is sensitive to assumptions about the life cycle, it may give robust estimates of the selection that maintains divergence in nature.

We also used a novel approach to estimate the strength of selection. Most studies of hybrid zones infer selection from cline widths which are determined by fitting descriptive cline models to allele frequency data (Fig. 3B). This approach is problematic. First, comparisons between single estimates of the width or centre can be misleading if they are made by fitting different models, and are given without support limits (which may be wide even when sample sizes are large). Moreover, the estimates of width can vary substantially depending upon the cline model that is fitted (Baird and Macholán, 2012). Indeed, the choice of model partly explains the differences in width between our study and other analyses of this zone which only fit a sigmoid cline (Bradley et al., 2025; Field et al., 2025). Second, the more complex descriptive models are not based on an explicit model of selection, and include impossible discontinuities in slope. Rather than relying on estimates of width, we inferred selection using a simulation framework that was underpinned by an explicit model of selection. This allowed us to include the full dispersal kernel together with multilocus selection. Although the diffusion approximation would be accurate in the limit of weak selection (assuming finite moments), here selection is moderate and long-distance dispersal is so strong that *σ*does not accurately describe dispersal in this system. By using the full dispersal kernel, we obtained estimates of selection that are quite different from those obtained using traditional theory, which substantially underestimated *s* for some loci (Table 3, Fig. S12).

Nevertheless, our framework has important limitations. First, it approximates the effects of dominance, epistasis and frequency dependence by assuming a linear change in selection coefficient across the hybrid zone, and does not include the additional LD that might be generated by epistasis; this might explain why the simulation fails to fully capture the observed cline shapes for some loci. We also assumed selection on diploids and equal haplotype frequencies in seed and pollen, but in reality, selection on flower colour is thought to act mainly through pollination and the actual life cycle depends on movements of both seed and pollen. Finally, the simulation framework ignores the effect of heterogeneous population density, which is known to shape patterns of isolation by distance in the snapdragon hybrid zone (Surendranadh et al., 2022)). Elaborating the model to include these factors may improve the fit between the observed and simulated data, yielding more precise estimates of selection in future.

Our simulation scheme allowed us to disentangle some potential causes of the stepped cline shapes. Specifically, we were able to separate the effects of long-range dispersal and LD on cline shape, showing that long-distance dispersal plays a much greater role. However, other factors may have contributed to the stepped shape. First, cline shape is determined primarily by the effective selection on each allele, which must be negative on one side and positive on the other: our basic model simply estimates these two parameters. However, dominance, epistasis and frequency-dependent selection will cause additional distortions in cline shape, but these are hard to disentangle from the strong effects of dispersal; we are skeptical that we could fit a more complex selection model. Second, our SNPs are only markers, linked to the causal alleles that are actually selected. Although a linked neutral locus will experience a barrier to gene flow due to its association with a selected one, it will inevitably recombine onto the alternative genetic background, allowing it to introgress across the cline. This introgression causes the cline to become stepped, with more pronounced tails of introgression observed for SNPs further along the chromosome from the selected site. As discussed above, future work will aim to explore this effect by jointly fitting the set of marker clines around each selected locus.

Finally, our study highlights the need for more nuanced interpretations of cline shape that go beyond explanations from traditional hybrid zone theory. The conventional interpretation of stepped clines is that they reflect a multilocus barrier to gene flow which reduces diffusive gene exchange across the cline centre. In our study, however, we found that stepped clines are instead primarily caused by long distance dispersal. Thus, we would substantially overestimate the strength of the genetic barrier to gene flow if we based this on the ratio between the size of the central step (spanning ∼1km), and the gradient in allele frequency on either side (*B* = Δ*p/p*^*′*^; Szymura and Barton (1986); Westram et al. *(2022))*. We show that reproductive isolation in snapdragons is due to reductions in two components of dispersal - namely diffusive gene flow and long-range dispersal. Further work is needed to deepen our understanding of how hybrid zones function as barriers to gene exchange, how those barriers can be estimated and to explore the effect of long-distance migrants on RI in other systems.

## 4 Methods

### 4.1 Sample collection and processing

The study site was visited annually from 2009 to 2019 during the main flowering season (late May-early August), aiming to sample every flowering individual. Within the study area, the same locations were re-visited several times over the period, to ensure that we captured plants that flowered at different times during the season. The sampling area encompasses a central region where the majority of plants are hybrids, and flanking regions with mainly parental types (Fig. 2C, D).

The geographic location of each plant was recorded with GeoXT handheld GPS units (Trimble, Sunnydale, CA, USA). Repeated visits to the sampled plants over the years have allowed us to quantify the mean position error of these GPS units as ±3.7 meters. We also collected leaf tissue for later genotyping, preserved by desiccating in silica gel. Additionally, we collected one flower to score the intensity and pattern of the yellow and magenta pigments.

### 4.2 Scoring of flower colour

Each flower was visually scored for magenta and yellow colouration based on the intensity and spread of the pigments (Fig. S2). The magenta scores ranged from 0.5 to 5, with a score of 0.5 showing no presence of magenta pigment to 5 showing intense magenta pigmentation throughout the corolla. Similarly, each flower was scored for yellow colouration, ranging from 0.5 (no yellow or yellow colour restricted to the bee entry point) to 3 (full yellow).

### 4.3 Development of SNPs and genotyping

A panel of 248 KASP SNPs spread throughout the genome was developed previously (see Methods in Ringbauer et al. (2018); Surendranadh et al. (2022); Field et al. (2025)). These include markers for divergent loci within or tightly linked to genes responsible for colour differences in snapdragons. All genotyping was conducted by LGC Genomics. Furthermore, error rates of these SNPs, estimated from 6000 plants that were genotyped more than once, were found to be low with an average probability 2 * 10^−4^ of misassignment to a different genotype (Field et al. in prep). We selected representative SNPs that are linked to known flower colour genes to understand the action of selection and dispersal on spatial genetic variation (Table 1). We had multiple linked loci to choose from, so we picked the one that showed the highest allele frequency difference between magenta and yellow ends, while also having low missing data.

### 4.4 Clustering individuals into demes

Rather than considering the individual position of each plant, we clustered neighboring plants into local demes. This was done for two reasons. First, the precise location of each sampled individual makes hardly any difference to the likelihood of a model in which allele frequencies change over a broad spatial scale but greatly increases the computational burden of cline-fitting. Second, we include fluctuations in the true allele frequency in our cline model, which are generated by the evolutionary process, as well as sampling error, to ensure that large demes are not given undue weight (see ‘Fitting clines to genetic data’).

Our clustering algorithm randomly selects the first individual, and then groups all individuals within the specified radius *δ* of the focal plant into a deme. It then moves on to the next available individual and repeats the process until all individuals have been grouped. We initially clustered individuals for a range of values of *δ*= {10, 15, 20, 25, 30, 40, 50, 75, 100, 120, 150, 175, 200, 250, 300m} and calculated the realized number and size of demes (Table. S1). From these, we chose a deme size based on two criteria. First, we checked if there were enough individuals in each deme to estimate local allele frequencies and LD. Second, we checked that *F*_*IS*_ was not inflated by the Wahlund effect (Wahlund 1928); (i.e., as *δ* increases, *F*_*IS*_ increases, due to isolation by distance). This was assessed by checking for departures from Hardy-Weinberg equilibrium for each deme size (see section ‘Deviations from HWE’ below). The spatial coordinates for each deme were calculated as the average from all plants within the deme.

### 4.5 Mapping onto a 1D transect

In Val de Ribes, the population of *A.m. majus* follows the south-facing slope of the valley, and so is essentially one-dimensional, with snapdragons primarily being found within 100m of two roughly parallel roads. We transform the data by mapping locations {*x, y*} onto a one-dimensional transect line. The distance along the transect is denoted by *z*. We established the transect line by fitting polynomials of degrees 1 to 8 to the deme positions, and minimising the mean square residuals (see SM S1). A polynomial of degree 6 was found to fit the data well (white curve in Fig. 2C and D), with most plants following the curve. We found little improvement from higher-order terms. For each deme, the point on the curve closest to the deme is found, and the new coordinate *z* is the distance to this point along the curve. The transect has an arbitrary starting point {410300, 4684000} and ranges from 0 to ∼30 km.

### 4.6 Fitting clines to genetic data

We fit three cline models to the frequency of the *pseudomajus* allele along the transect for all clinal SNPs. The first model (sigmoid) has allele frequencies ranging from 0 to 1, as predicted for selection against heterozygotes (Bazykin, 1969), symmetric epistasis (Bazykin, 1973) or frequency-dependent selection with no dominance, 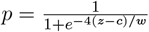 where *c* is the cline centre and *w* is the cline width (Fig. 1A); other single-locus models give a similar shape (Barton and Gale, 1993). We also include random variation in allele frequency around the cline, measured by *F*_*ST*_; sampled frequencies are binomially distributed around the underlying frequency, which follows a Beta distribution with variance *pqF*_*ST*_ (Slatkin and Barton, 1989). Thus, we have three parameters {*c, w, F*_*ST*_}.

For the second model (sigmoid with polymorphism), we adjust the sigmoid curve by allowing for polymorphism on either side: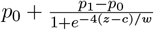. This model has 5 parameters {*c, w, p*_0_, *p*_1_, *F*_*ST*_}.

For the third model (stepped), we splice exponential functions onto the left and right sides, 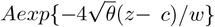 (Szymura and Barton (1986, 1991), Fig. 1B). This provides a model of a stepped cline with a sigmoid shape in the centre and long exponential tails of introgression on either side. The parameter *θ* gives the rate of decay of the tail, slower than predicted by the simple sigmoid model (i.e. *θ* ¡1). The constant *A* is defined by the strength of the barrier to gene flow, defined by *B* = Δ*p/p*^*′*^, where Δ*p* = *p*_1_ − *p*_0_ is the difference in allele frequencies at the transition from sigmoid to exponential and 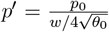 is the gradient in allele frequency at the edge of the central segment (see electronic SM code for details). Introgression may be symmetric or asymmetric, and so we estimate {*θ, B*} separately on either side. Thus, we have seven parameters *c, w, θ*_0_, *B*_0_, *θ*_1_, *B*_1_, *F*_*ST*_.

The parameters are estimated using the Metropolis Hastings algorithm (Metropolis et al., 1953; Szymura and Barton, 1986), which generates a random walk that follows the likelihood: random changes are made to the parameter set and the changes are accepted if they increase the likelihood or rejected otherwise. More formally, we start with an initial parameter set and its likelihood *L*. Uniformly random changes [−*δ, δ*] are made to each parameter sequentially and the new likelihood *L*^*′*^is calculated. If *L*^*′*^*/L* is greater than a uniform random number between 0 and 1, the change is accepted and parameters and likelihood are reset to the new values. Further, we set *δ* to *λδ*, where *λ* =1.01. If *L*^*′*^*/L* is less than the random number, we reject the change, but set *δ* to *δ/λ*. With enough trials, this will generate a distribution proportional to *L*. The size of changes, *δ*, is tuned for efficiency, ensuring that similar numbers of changes are accepted as rejected. Cycling through each set of parameters 3000 times assured convergence.

For each locus, this fitting process was conducted separately for each model. The best fit of the three cline models was determined for each locus using likelihood ratio tests (Kawakami et al., 2009). For the best model, the maximum likelihood estimates (MLE) of the parameters were used to obtain the cline shape. The Metropolis Hastings algorithm not only gives the MLE but also provides the marginal likelihoods of parameters.

### 4.7 Comparing clines over time

The stepped cline model was employed to fit clines for *ROS* at various time points, since this model gave the best fit when considering all years collectively. 11 years of data are separated into three time points; 2009 to 2012, 2013 to 2015 and 2016 to 2019 with 7690, 12409 and 7384 individuals respectively. Pooling individuals from 3 to 4 years into a single time point is a valid assumption as *A.m. majus* has overlapping generations and average dormancy of 3-4 years (as observed from the pedigree trios in Field et al. in prep.).

### 4.8 Cline fit for the phenotypic data

Magenta and yellow colour scores for each individual were rescaled so that they fell between 0 (minimum) and 1 (maximum). We then calculated the mean colour score for each deme which ranged from 0 to 0.889 for magenta and 0 to 1 for yellow. We fit the same three cline models to these mean values to identify the best-fitting model and MLE of each parameter, as described above for the genetic data (see section ‘Fitting clines to genetic data’).

### 4.9 Deviations from Hardy Weinberg equilibrium

The deficit of heterozygotes relative to the Hardy-Weinberg expectation, *F*_*IS*_, is calculated for each deme, as one minus the ratio of observed and expected number of heterozygotes for each unlinked clinal locus at the clustering scale of *δ* =25m. This was done by first selecting 337 of 999 demes with at least 10 individuals, and then dividing the transect into 12 bins such that each bin contains a roughly equal number of demes (see Table S4). The mean and standard deviation of *F*_*IS*_ were computed for each bin. A null distribution of *F*_*IS*_ was also generated for each locus and deme by making 1000 replicate draws of diploid genotypes, fixing allele counts as observed, but combining alleles at random. The observed and the null distribution were then compared to check for demes (and bins) that showed significant deviations from Hardy Weinberg equilibrium. Significance was based on the p-value for each deme, i.e. the fraction of shuffled replicates with *F*_*IS*_ less than the observed if the latter is less than 0 and greater than observed otherwise.

### 4.10 Linkage disequilibrium based on hybrid index

Linkage disequilibrium (LD) between two unlinked loci was estimated from the variance in hybrid index (HI) as 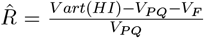, where *V*_*P Q*_ is the variance due to heterozygosity and *V*_*F*_ is the variance due to heterozygote deficit. Since missing data could be correlated across loci and generate a positive bias in the estimator, 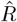 was measured based on pairs of loci. Individuals with missing data at either of the two loci in consideration were deleted and 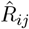 was estimated based on HI calculated from the 2 loci. 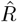 is then the average of 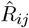 for *i j*. 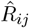 was first estimated for each locus pair for the whole population to check for significant associations between loci. Next, we looked at how average associations between loci change across the geographic range. 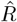 was found for the 12 bins as above (see section ‘Deviations from Hardy Weinberg equilibrium’) by considering 337 demes with at least 10 individuals. In each case, a null distribution was generated by shuffling genotypes across loci for 1000 replicates, thereby preserving HW deviations but generating linkage equilibrium. A p-value was obtained for each deme by finding the fraction of shuffled replicates that gave values less than observed if the latter is less than 0 and greater than observed otherwise.

### 4.11 Estimating long-range dispersal for *A.m. majus*

2342 pedigree trios inferred by Field et al. (in prep.) give estimates of pollen and seed dispersal, which extend out to ∼1km. he pollen dispersal is close to a log-Gaussian distribution for distances up to 600 m that can be estimated reliably from the trios (Fig. 5). We used this log-Gaussian distribution for pollen dispersal, *extrapolating it out* to longer distances: 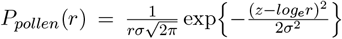, where the MLE of mean (*z*) and standard deviation (*σ*) of *log*_*e*_*r* were 2.88 and 1.64 respectively (corresponding to a geometric mean pollen dispersal distance of *e*^2.88^ = 17.8m).

However, longer-range dispersal will be underestimated from the pedigree trios, because parents are increasingly likely to be outside our study area. We do not make a detailed analysis of this bias, but instead, only consider seed dispersal up to 300m, which is typically much less than the distance to the nearest populations outside our study area. Longer-range seed dispersal beyond 300m is estimated from individuals that have genotypes most likely to come from the opposite side of the hybrid zone. We treat the yellow and magenta flanks separately, and now work in one dimension, since that better describes the transect over large scales.

The actual population has a complex spatial distribution, concentrated around two main roads, but spreading out to either side. Estimates based on the pedigree trios (Fig. 5A) approximate shortrange dispersal (*<* 300*m*) as two-dimensional, described by a radial distribution. The one-dimensional distribution along some axis (which required for fitting clines) is derived by convolving with 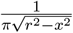, which is the probability of moving in a random direction, given *r*. We can then derive the distribution of net short-range dispersal by using the fact that a gene has an equal chance of coming from the mother (via seed dispersal), or the father (via seed plus pollen dispersal). Therefore, the net distribution is an equal mixture of the seed dispersal, and the convolution of seed with pollen dispersal.

We estimate longer-range dispersal (up to several km) from individuals that have genotypes typical of the opposite side of the hybrid zone, based on the 5 unlinked SNP; these individuals are presumed to have arrived by seed dispersal. Table S8 assigns individuals as immediate dispersers, F1, backcross, …, based on the observed allele frequencies in the flanks, and assuming Hardy-Weinberg and linkage equilibrium (HWLE; see S3); this shows that a few percent of the populations in the flanks are likely to have arrived by seed dispersal in the previous generation. However, we estimate long-range dispersal using a more sophisticated method, based on the likelihood of the mother’s location, and assuming exponential tails. (We have little power to distinguish alternative kernels in the tails, so this choice is arbitrary). The probability *P* (*X, z*) of finding an individual with genotype *X* at location *z* is: 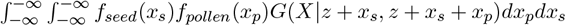. This is an integral over the probability *f*_*seed*_(*x*_*s*_) that the mother was *x*_*s*_ away, times the probability *f*_*pollen*_(*x*_*p*_) that the father was *x*_*p*_ away from the mother, times the probability *G* that the offspring has genotype *X*, given the allele frequencies at the parents’ locations, assuming HWLE; this is calculated as a product of allele frequencies across loci, with factors {*q*_*y*_*q*_*z*_, *p*_*y*_*q*_*z*_ + *p*_*z*_*q*_*y*_, *p*_*y*_*p*_*z*_}, where *p*_*y*_ and *p*_*z*_ refer to the allele frequencies at locations *y* and *z* respectively predicted from the cline fits at that locus (Fig. S7), and *q* = 1 − *p* (see S3 for more details). *f*_*seed*_ is the seed dispersal distribution, which takes the form estimated directly from the trios up to 300m, and is then spliced onto 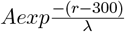 for distances greater than *x*= 300m. The log likelihood for *λ* in either flank is estimated by summing *log*(*P* (*X, z*) across all individuals in the dataset; *A* is determined by splicing at 300m.

#### 4.1.2 Estimating strength of selection from simulated clines

##### Simulation framework

We consider a deterministic stepping stone model with 45 unevenly spaced demes that range from -13.5 to +13.5 km. The demes are located at {…, −*δϕ*^2^, −*δϕ*, 0, *δϕ, δϕ*^2^, …} where *δ* =50 and *ϕ* =1.2. This spacing was chosen so that demes span the length of the one-dimensional transect, but are concentrated near the centre, where allele frequencies change most rapidly. We start with a sigmoid cline of width 4 km centreed at 0, which gives the initial frequencies of 2^*l*^ haplotypes at each deme, assuming *l* biallelic loci under linkage equilibrium.

We use the estimated dispersal distribution to simulate clines along a one-dimensional transect. Each deme is centred at a point, and extends half-way towards its neighbours on either side; the fraction that move is given by integrating over the span of the deme. We assume uniform density, so that the number in each deme is proportional to its span. The backwards migration matrix is calculated from the CDF of net dispersal, by calculating the fraction of genes in each deme that came from every other deme in the previous generation. Because we estimated different exponential tails on either sides, we interpolate their distance scale using a logistic function, such that at distance *z* along the transect from the centre, 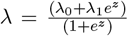. Asymmetric dispersal causes the clines to move. To overcome the computational burden of recalculating the migration matrix, we fix the locations of demes and shift the clines to be centred at 0 every 10 generations. The cline centre is defined as the location at which the gradient of the interpolated cline is steepest.

The simulation follows the haplotype frequencies at each deme for 500 generations; random union of gametes is followed by selection on diploid genotype, meiosis and migration of haploids. Note that the simulations only follow haplotype frequencies under HW equilibrium, assuming the same allele frequencies in pollen and ovule. We simulated clines at a single locus with heterozygote disadvantage and Gaussian dispersal to check that simulations gave results consistent with theory.

##### Estimating selection coefficients

This model was used to simulate clines, either considering a single locus or 5 unlinked loci. As above (see ‘Fitting clines at genetic loci’), we assume that observed allele frequencies are sampled binomially from an underlying Beta distribution, with variance *Fpq*. When we fit single loci, we assume symmetric selection *s*(fitnesses 1+*s*: 1: 1+*s*), and estimate *s, c* (cline centre), *F* (standardised variance), giving 5*3=15 parameters across 5 loci. For multilocus clines, we allowed asymmetric selection at each locus (fitnesses 1 + *s*_*left*_: 1: 1 + *s*_*right*_), multiplicative across loci; we assume a single value of *c* and *F* for all loci, giving a total of 2+5*2=12 parameters.

For single-locus clines, we first maximise likelihood estimates of *c* and *F* for given *s*, and then find the most likely *s*; this avoids unnecessary simulation. With multiple loci, the MLE of parameters at each locus depends on the allele frequencies at all other loci. We start with the MLE of *s* from the single locus fits, and then follow the single locus procedure for each locus in turn, finding a new set of selection coefficients, now allowing asymmetry (*s*_*left*_, *s*_*right*_). This procedure is repeated until estimates converge.

## Conflict of Interest

The authors declare no conflict of interest.

## Acknowledgements

We thank Enrico Coen, Desmond Bradley, and other members of the Coen Group at the John Innes Centre for their extensive contributions to the fieldwork, development of genetic resources and detailed insights about the genetic basis of colour upon which this study depends. We thank all the interns and field managers, including Eva Cereghetti, Beatriz Pablo Carmona, Mariona Vinyeta, Helena Ramirez, and all volunteers who have helped us with field work and processing over the years. We thank Eva Salmerón Mateu for her help with logistics at the field station in El Serrat, Planoles. We also thank all the members of the Barton Group, especially Tom Ellis, Harald Ringbauer, Melinda Pickup, Carina Baskett, Maria Melo Hurtado for supporting the fieldwork and for feedback on this work.

## Funding

This work was funded by Austrian Science Fund FWF (grant P32166) and ERC grant (101055327).

## Author Contributions

Team led by Enrico Coen and Nicholas H. Barton established the intensive sampling scheme which was initially funded by a BBSRC grant awarded to EC and later funded by NB. All authors contributed to the fieldwork. PS, DF and NB conceived the idea, DF built the data management pipeline and generated the SNP panel, PS and NB performed the analyses. PS, SS and NB wrote the manuscript. All authors contributed to editing the manuscript.

## Supplementary Materials

## Supplementary Text

### S1: Finding the 1D polynomial transect

To fit clines, we converted the 2D spatial positions of the demes into 1D by fitting a polynomial transect of degree *n, y* = *a*_0_ + *a*_1_*x* + *a*_2_*x*^2^ + … + *a*_*n*_*x*^*n*^, where x is the E-W position. We fit polynomials of different degrees to the data, calculating the sum of squared distances from the line of demes (defined at 25m scale). This sum decreases with the degree of the polynomial, but the decrease flattens out beyond degree 6 (SMxx). For each deme, we find the point on the curve that minimises the distance between the deme and the curve. The distance of the point along the curve was obtained as 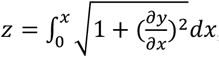, where *x* is the position of the deme on the curve.

### S2: *F*_*IS*_ at different clustering radii

To examine deviations from Hardy Weinberg equilibrium and choose a spatial scale that minimises departures from random mating, the maximum likelihood estimate (MLE) of *F*_*IS*_ is found at different clustering scales for the clinal SNPs (excluding *EL*, which is linked with *ROS*) and the 91 non-clinal SNPs (as in Surendranadh et al. 2022), assuming a uniform value across all demes. *F*_*IS*_ depends on the allele frequencies (*p,q*) and is bounded by max[{- *p/q, q/p*},1]. *F*_*IS*_ increases with deme size due to the Wahlund effect, with similar slopes for both clinal and non-clinal loci, as expected from isolation by distance. Heterozygote deficit was greater for clinal SNPs when compared to the mean from non-clinal SNPs (coloured vs black curves in Fig. S3). The magnitude of *F*_*IS*_ differs between clinal loci and is somewhat consistent with the difference in cline width; *ROS* shows the largest heterozygote deficit, and *RUB* together with the non-clinal SNP, the smallest. We see some *F*_*IS*_ over small scales, which may be due to small-scale local fluctuations at loci with steeper clines, or to assortative pollination for flower colouration. However, at the 25m scale, *F*_*IS*_ is close to 0 for all loci except *ROS* (see Fig. S3). Additionally, multilocus heterozygosity from 91 non-clinal SNPs was consistent with matings from parents 10m apart (see Fig. S1, Surendranadh et al. 2022). Taken together, 25m is a reasonable choice for the clustering scale.

### S3: Evidence of improbable individuals in the flanks

For each individual in the yellow and magenta flank, the full genotypes are examined at five unlinked loci, to identify recent migrants and their offspring. The individuals are assumed to either come from a local population (i.e deme in which the individual belongs), which is assumed to be well-mixed, or from a well-mixed distant source (i.e. the magenta flank if the focal individual comes from the yellow flank and vice versa) or are F1 or backcross hybrids between these sources.

The probability *P*_*genotype*_ of a diploid genotype in a native population that is in Hardy-Weinberg proportions and linkage equilibrium (HWLE) is the product of allele frequencies across loci, with factors {*q*_*i*_^2^, 2*p*_*i*_*q*_*i*_, *p*_*i*_^2^} according to diploid genotype (denoted X_i_ = 0,1 or 2). We estimate log (*P*_*genotype*_) as a sum over loci, and for missing values, add the average*q*_*i*_^2^ *log*(*q*_*i*_^2^) + 2*p*_*i*_*q*_*i*_ *log*(*p*_*i*_*q*_*i*_) + *p*_*i*_^2^ *log*(*p*_*i*_^2^) (4.1% of values are missing at these loci). This extends in a simple way to F1s and backcross hybrids derived from a cross with a source population with allele frequencies *pj*. The frequencies of the three diploid genotypes are now {*q*_*i*_*q*_*j*_, *p*_*i*_*q*_*j*_ + *q*_*i*_*p*_*j*_ + *p*_*i*_*p*_*j*_}, and the net probability is again a product over loci. As before, we calculate log(*P*), add the expected contribution when there is a missing value, and account for the error rates in this procedure. Note that this algorithm does capture the linkage disequilibrium generated by admixture, even though it is based on a product of factors across loci, and extends to individuals from any two locations.

For each flank, we first identified individuals with *P*_*genotype*_ < 2 *\** 10^−4^ of coming from their location, with *P*_*genotype*_ calculated as described above. Here, the allele frequency is based on the best estimate from the cline fitting for that location (*see Cline fitting for the genetic data*). For the ‘improbable individuals’ that are unlikely to come from the local populations, we generated the probability that it came from the opposite flank, or is an F1 or backcross derived from that source. The allele frequency of the source population is calculated directly from the genotypes of all individuals in that flank. Table S8 shows the improbable individuals from the yellow and magenta flanks.

## Supplementary Figures

**Fig S1:**
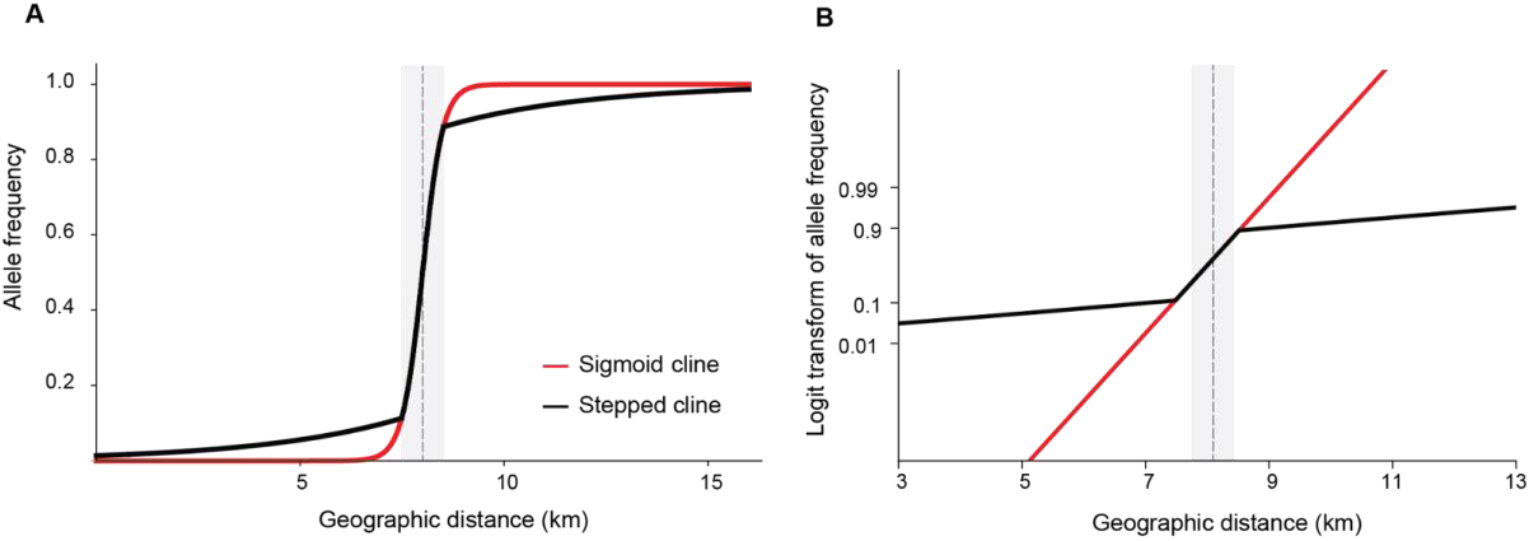
Sigmoid (red) and stepped (black) clines of the same width (1 km) and centre (8 km) plotted on normal scale (A) and logit transformed (B). The dotted line represents the cline centre and the grey bar denotes the cline width. (A) A stepped cline has a sigmoid shape in the centre and overlaps with the sigmoid cline of the same width in the centre. (B) The logit transformation plots log(*p/q*), where *p* is the allele frequency and *q= 1-p*. Here, the sigmoid shape appears as a straight line and the step is easily distinguishable.

**Fig S2:**
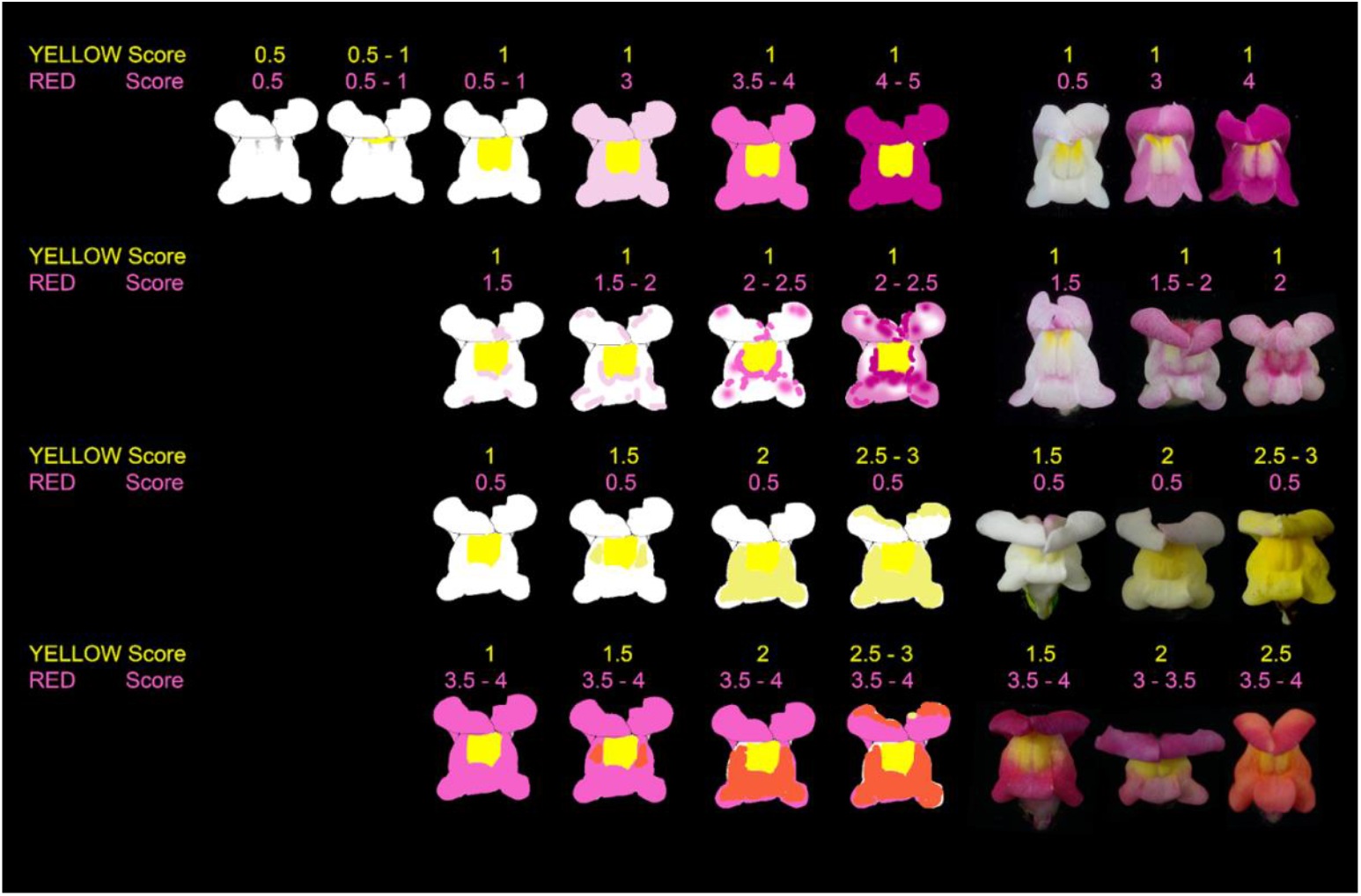
Flower scoring guide used to score the magenta (indicated as ‘Red’ in the figure) and yellow flower colouration. For each flower, magenta and yellow flower scores range from 0.5 to 5 or 3 depending on the intensity of the pigmentation and how it spreads across the flower.

**Fig S3:**
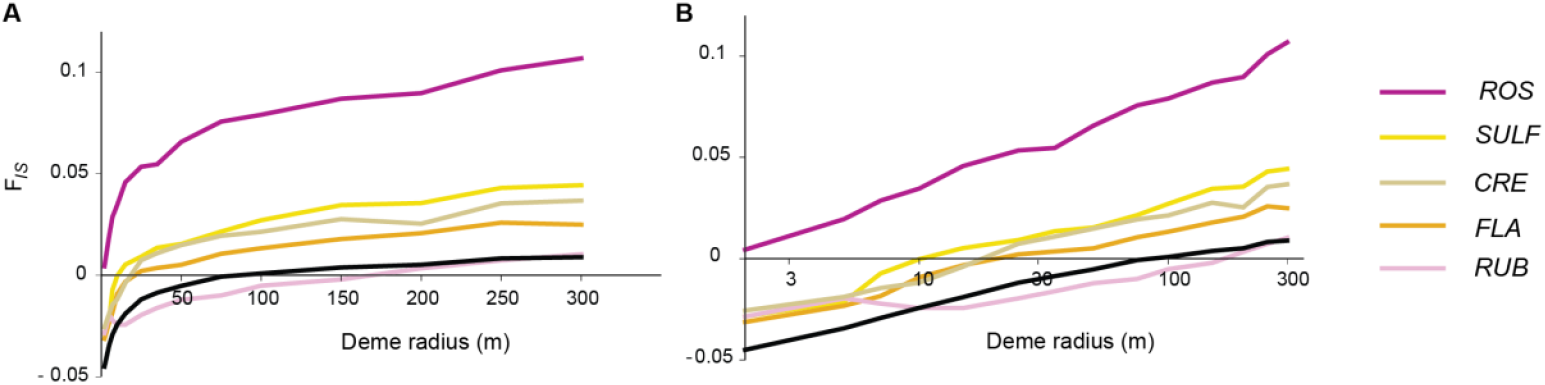
Mean heterozygote deficit (*F*_*IS*_) across 999 demes for different deme radii shown for each clinal loci on the normal scale (A) and (B) log scale. The mean *F*_*IS*_ from 91 non-clinal SNPs from Surendranadh et. al 2022 is shown in black for comparison.

**Fig S4:**
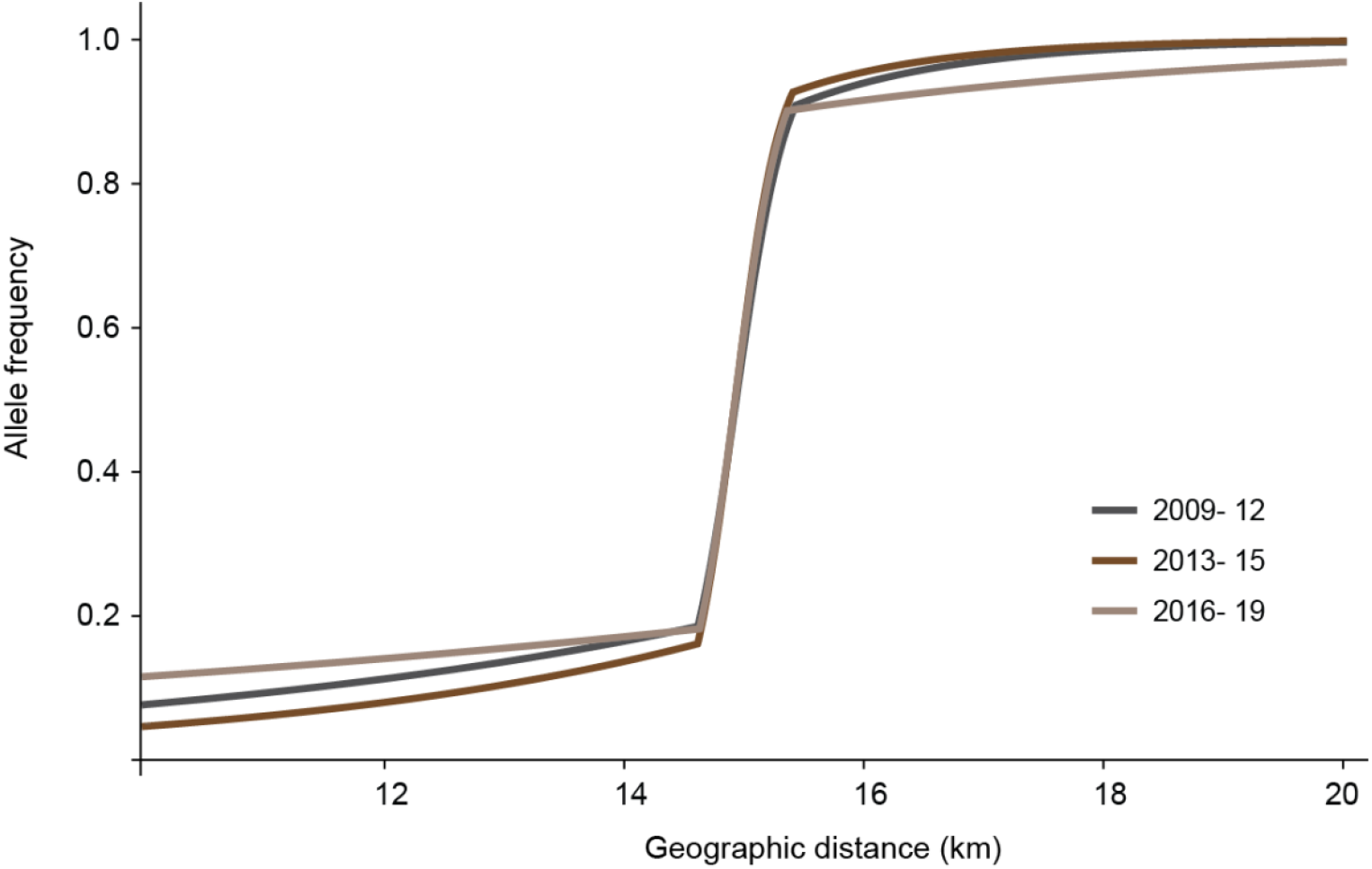
Temporal clines in *ROS*: clines are fitted for *ROS* across 3 time points 2009 to 2012, 2013 to 2015 and 2016 to 2019 shown as allele frequency plotted across a smaller range from 10km to 20 km. The central cline shape is consistent across different time points with similar cline centres and cline widths (see Table S3). However, the cline shape in the right and left tails differ due to differences in the polymorphism across years.

**Fig S5:**
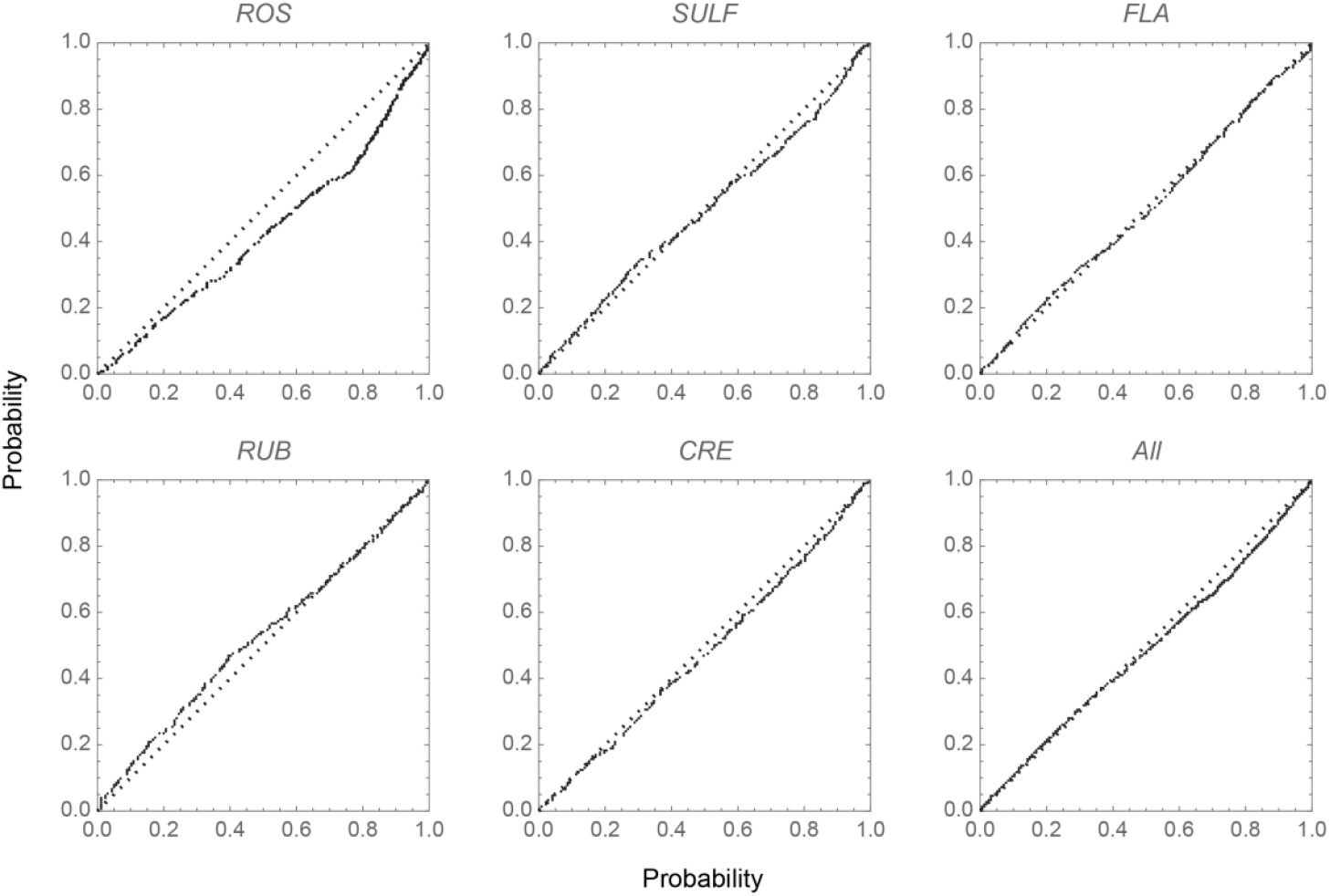
Q-Q plot showing observed and expected *F*_*IS*_ for each loci and mean *F*_*IS*_ from all loci. Each plot compares the observed distribution of *F*_*IS*_ from each deme to the null distribution obtained by shuffling within each deme.

**Fig S6:**
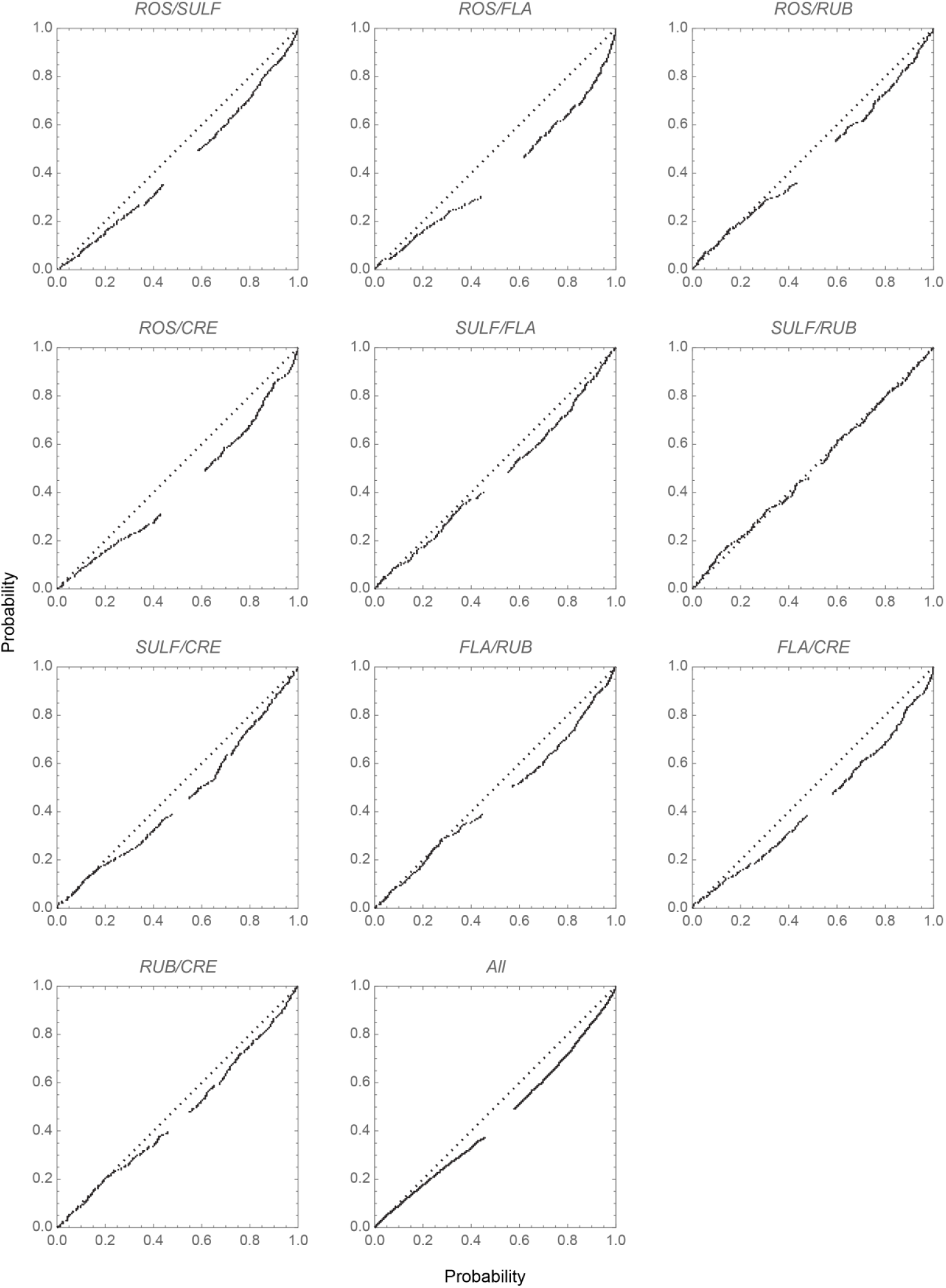
Q-Q plot showing observed and expected 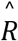 for each locus pair and mean 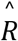 from all pairs of loci. Each plot compares the observed distribution of 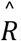 from each deme to the null distribution obtained by shuffling within each deme

**Fig S7:**
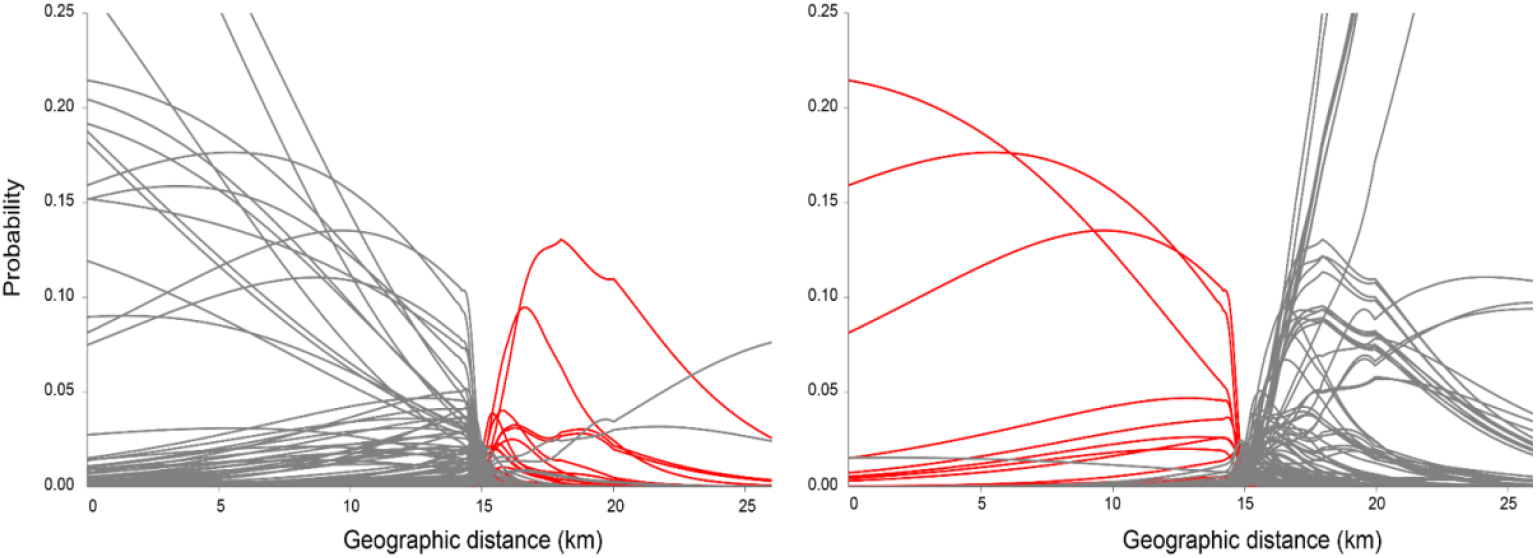
Marginal probability of finding the mother of an individual at a given distance plotted along the transect. Each curve represents an individual in the yellow flank (left panel) and magenta flank (right panel) respectively. Grey curves denote individuals in the flanks likely to come from a mother on the same side. Red curves denote individuals in the flanks likely to come from a mother on the opposite flank.

**Fig S8:**
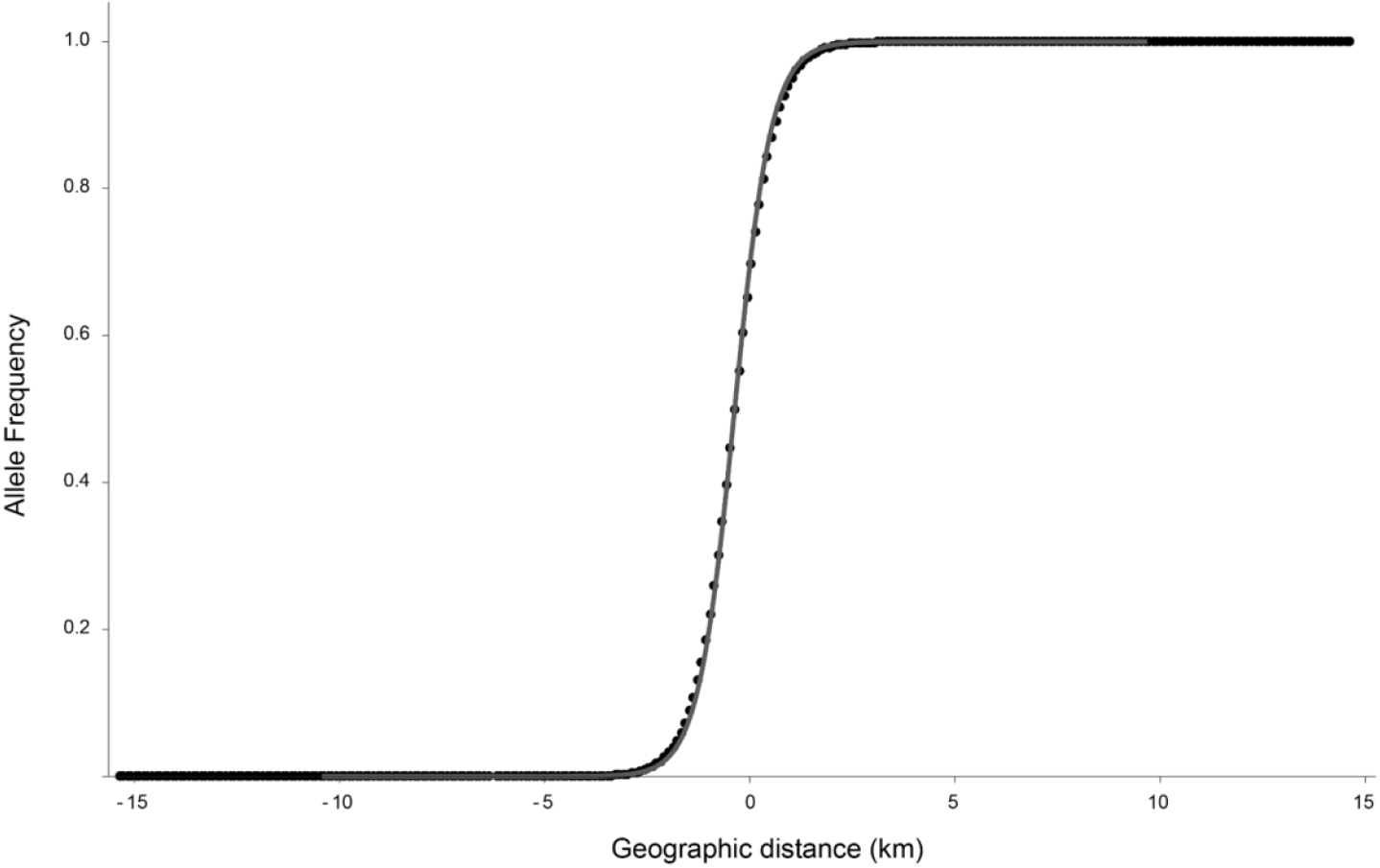
Allele frequencies versus geographic distance (km) from a single locus cline simulation considering Gaussian dispersal with *σ* = 200m and heterozygote disadvantage with *s* = 0.1. The black dots denote the simulated allele frequencies in the demes and the curve represents the allele frequencies from the expected sigmoid cline of width 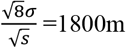.

**Fig S9:**
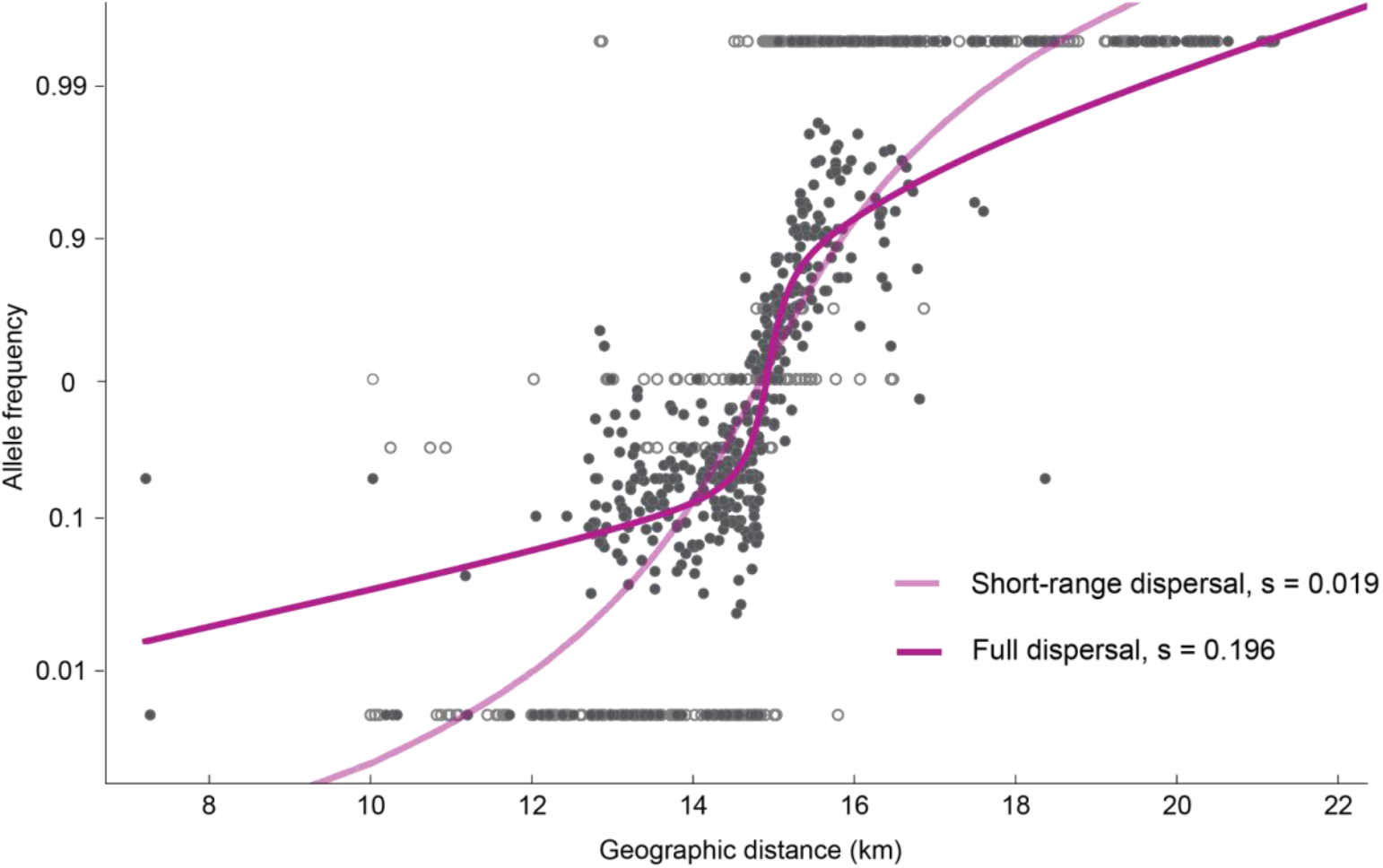
Simulated clines at *ROS* plotted on a logit scale (i.e allele frequency *p* is transformed into *p/q* and shown on a log scale). ML estimates of selection are inferred from two simulations, one considering short-range dispersal (i.e inferred only from the trios in Field et al. in prep.) where *s* = 0.019 (in light magenta) and with the full dispersal kernel where *s* = 0.196 (in dark magenta). The former fails to produce a strong step as observed. Simulated cline with the inferred dispersal generates a stepped shape and captures patterns of allele frequency in the tails.

**Fig S10:**
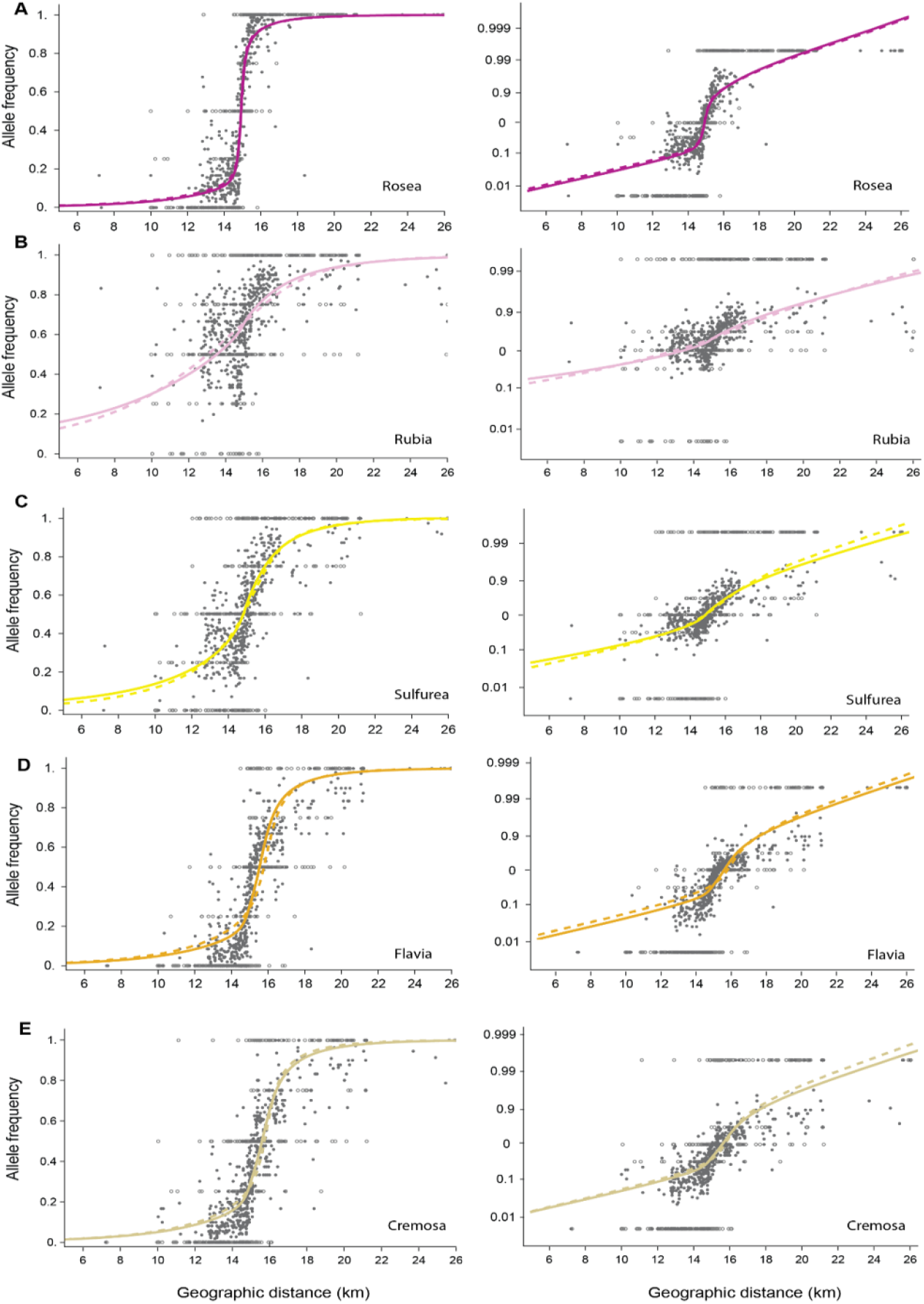
Best fit cline shapes from single locus simulations (dashed) and multilocus simulations with asymmetric selection (solid) at each of the 5 unlinked loci. The left column plots it in a normal scale while the right shows the same in the logit transformed scale. The corresponding selection estimates are shown in Table 3 of the main text.

**Fig S11:**
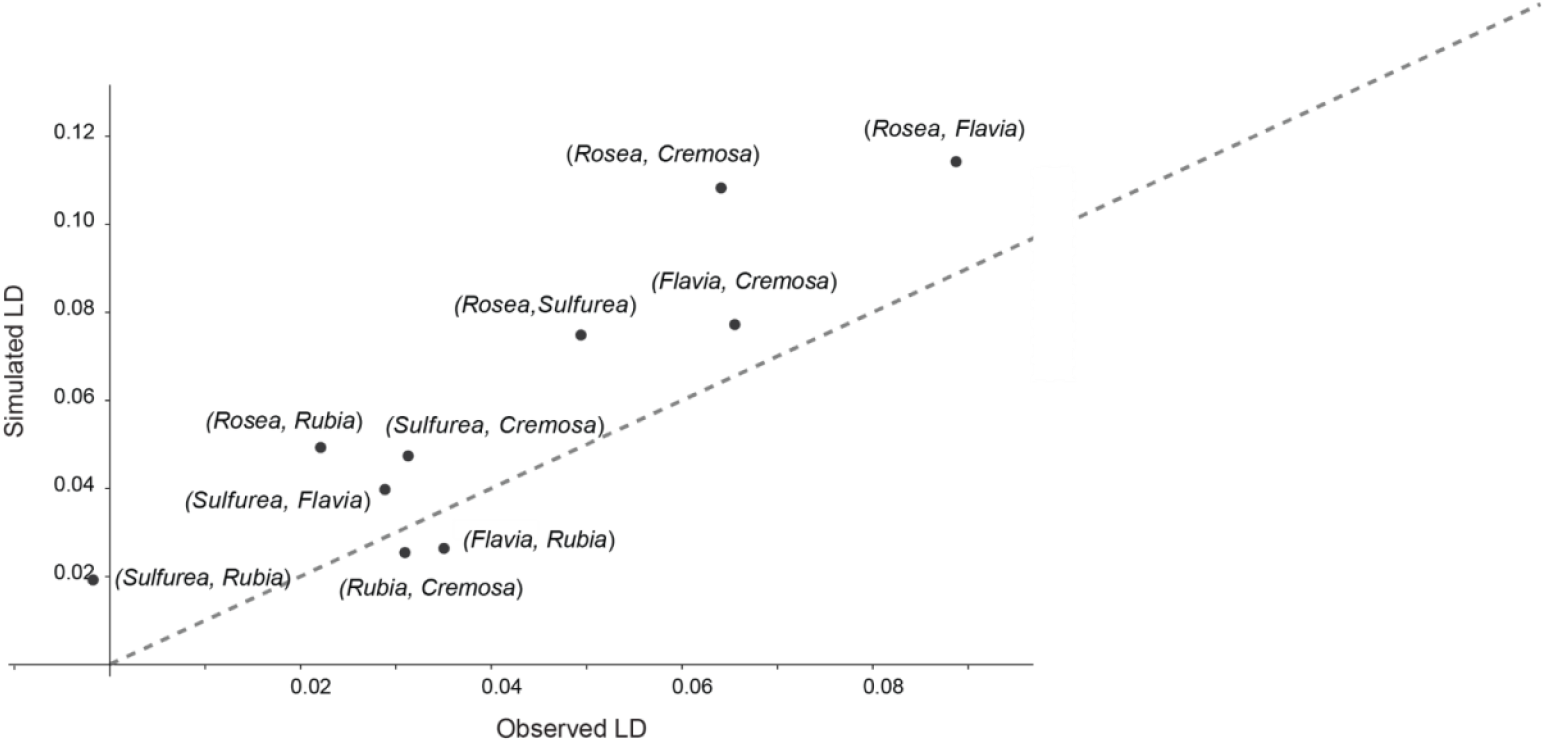
Comparison of LD between 10 pairs of unlinked loci from multilocus simulations with the inferred dispersal vs that observed from the data.

**Fig S12:**
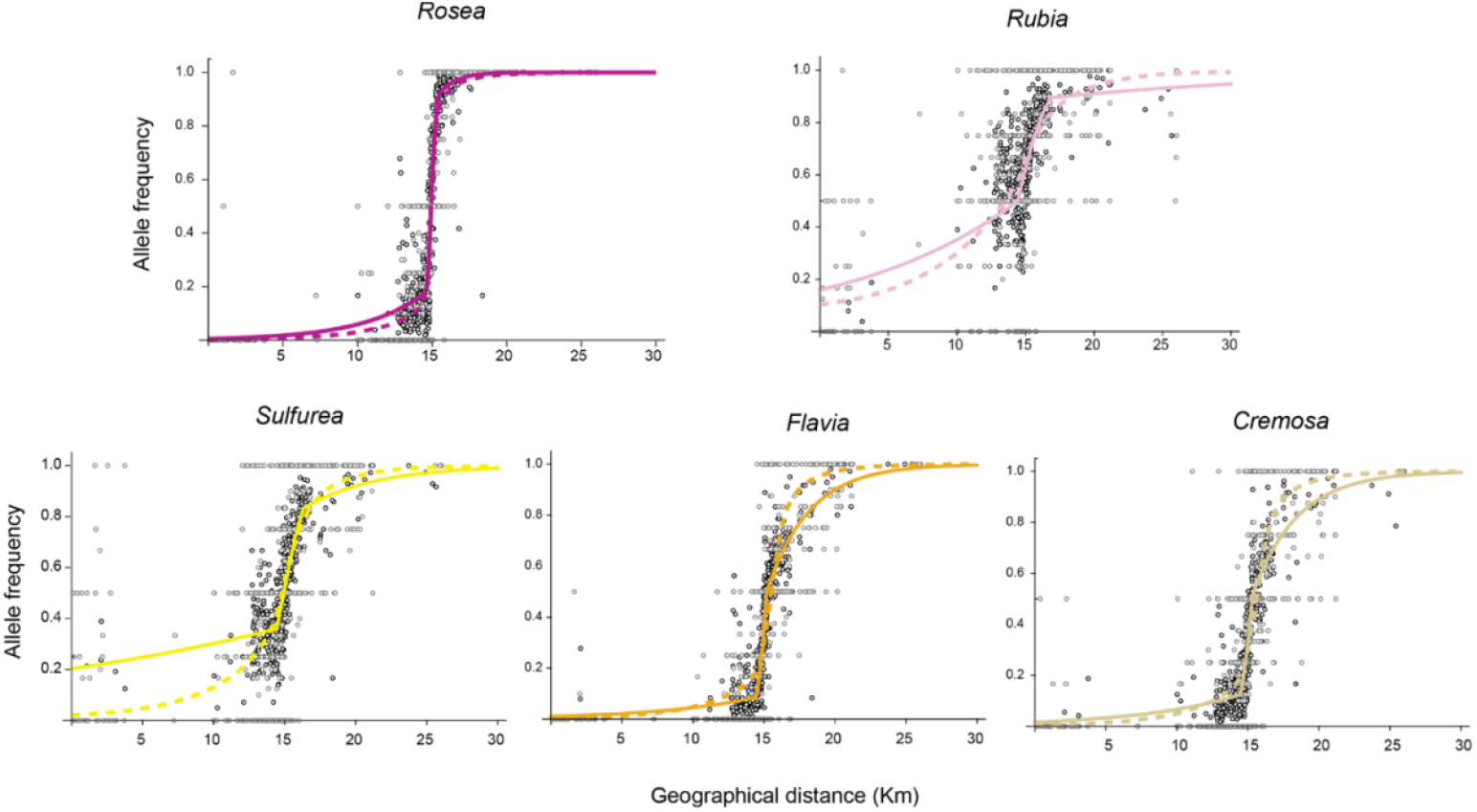
Best fit cline shapes from multilocus simulations with asymmetric selection (dashed) vs from descriptive cline fitting (solid) at each of the 5 unlinked loci. The corresponding selection estimates are shown in Table 3 of the main text.

**Fig S13:**
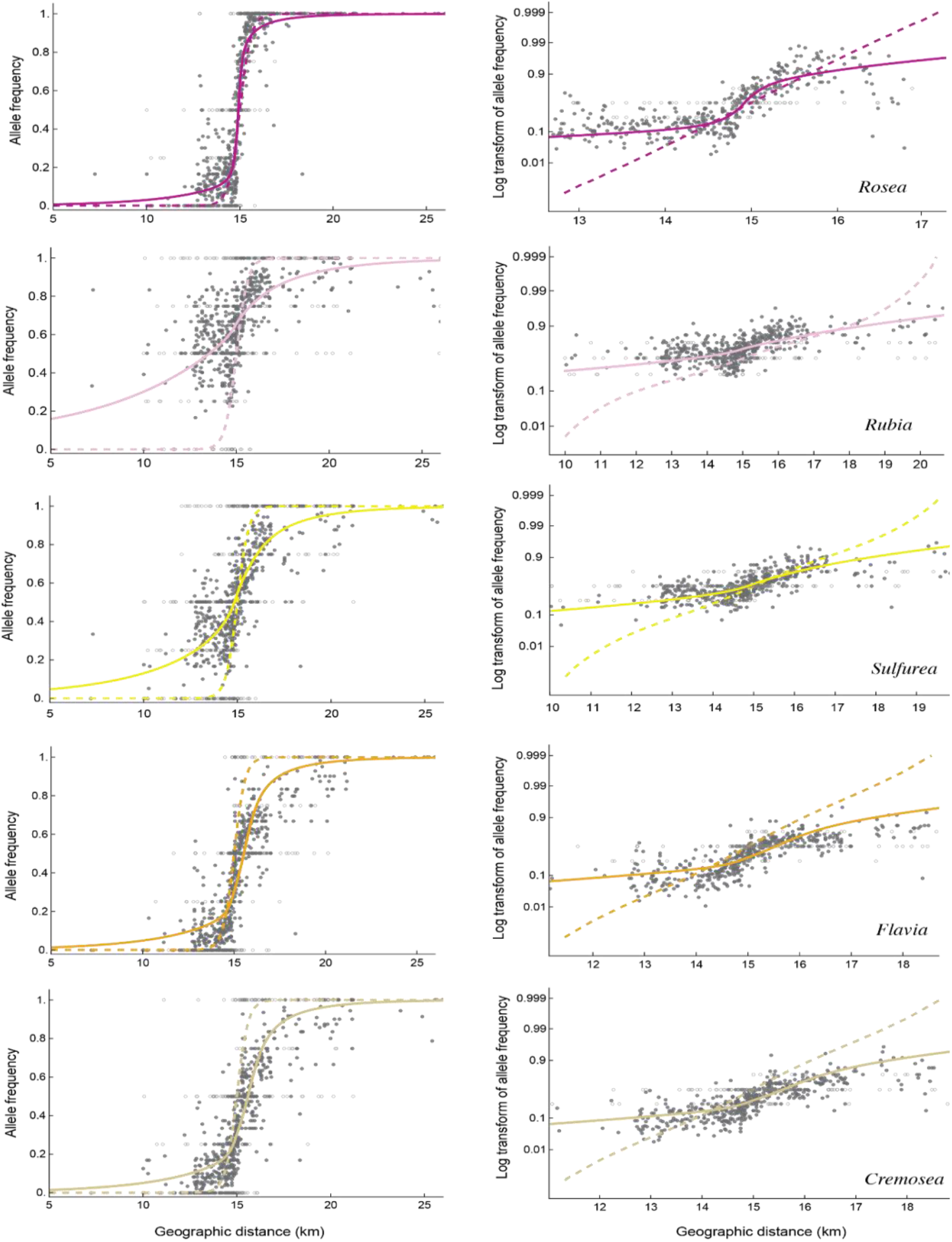
Best fit cline shapes from multilocus simulations with inferred long-range dispersal (solid) vs Gaussian dispersal with *σ* =161m (dashed) at each of the 5 unlinked loci. The left column plots it in a normal scale while the right shows the same in the logit transformed scale. The latter shows that multilocus simulation with Gaussian dispersal does not produce stepped clines, suggesting weak LD at the cline centre.

**Fig. S14:**
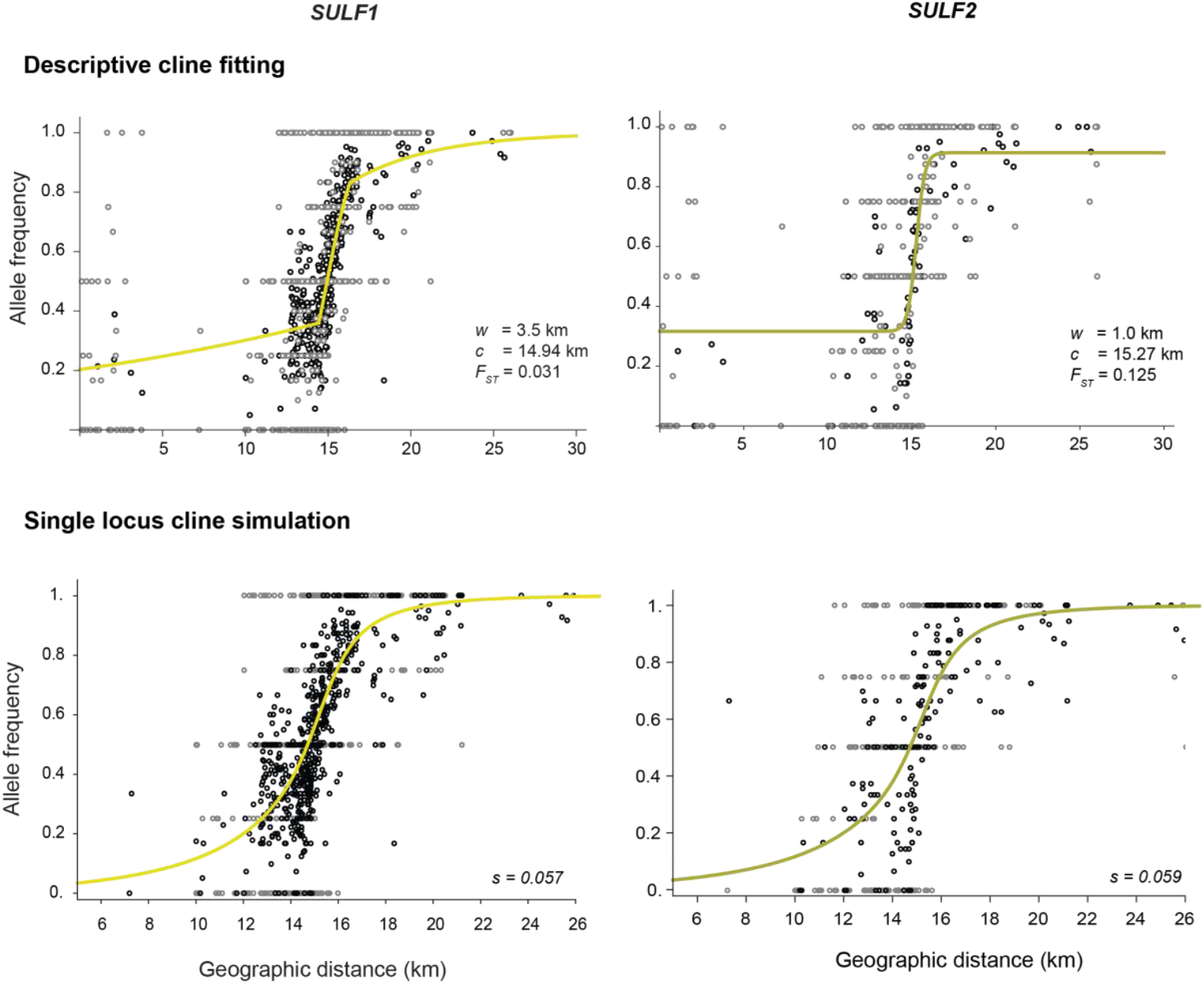
Comparison of cline shapes between two *SULF* SNP markers: *SULF1* analysed here vs *SULF2* analysed in Bradley et al. 2025. The cline width estimated for *SULF2* matches the prediction from Bradley et al. 2025, but is much narrower than *SULF1*. However, the strength of selection estimated from single locus cline simulations is comparable across both markers.

## Supplementary Table

**Table S1:**
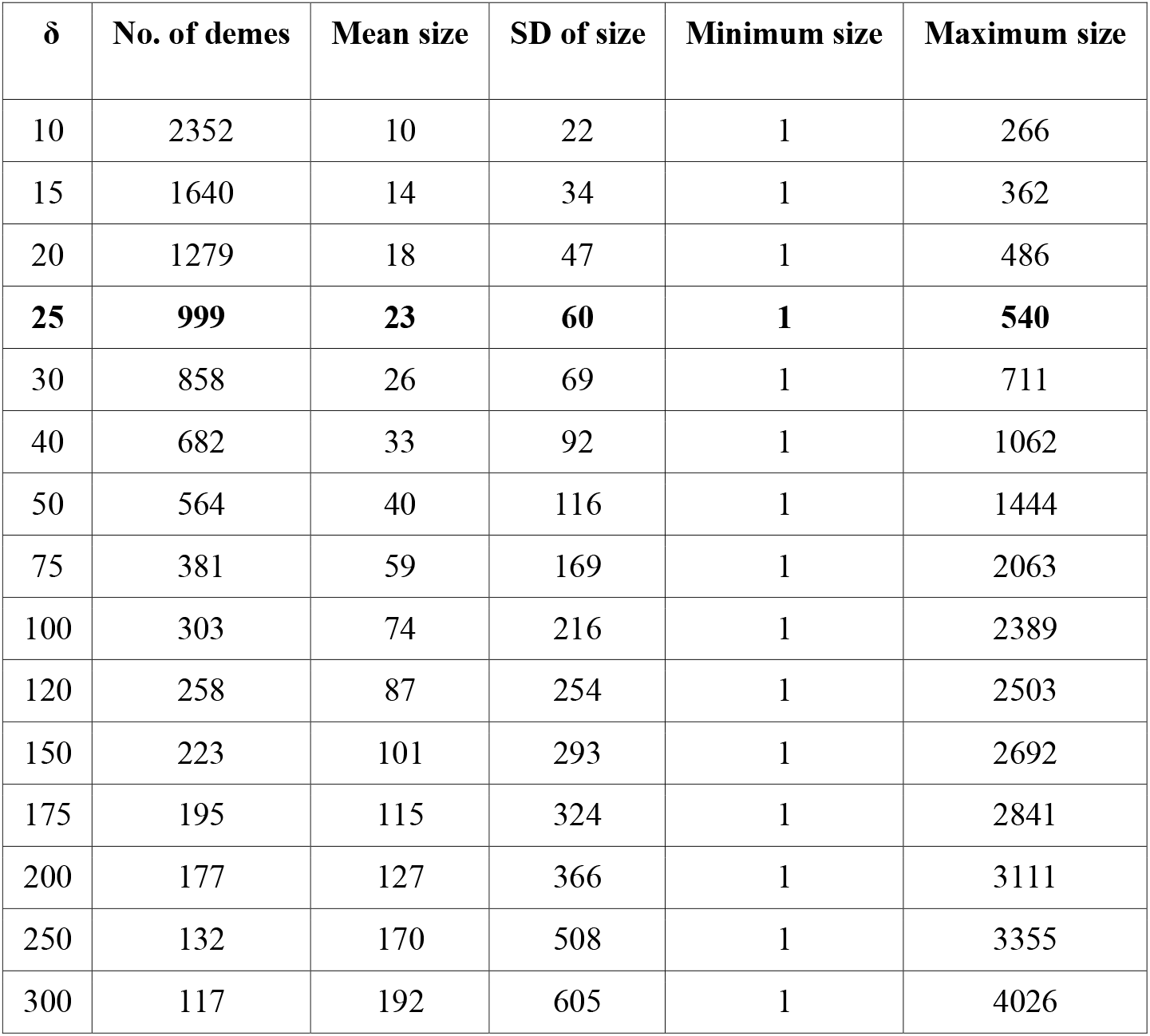
Number of demes, mean, standard deviation, minimum and maximum size of the demes corresponding to each cluster radius *δ*. The size of a deme denotes the number of individuals within a deme. The clustering radius of 25m was used throughout the analysis, giving 999 demes with an average of 23 individuals in each deme.

**Table S2:**
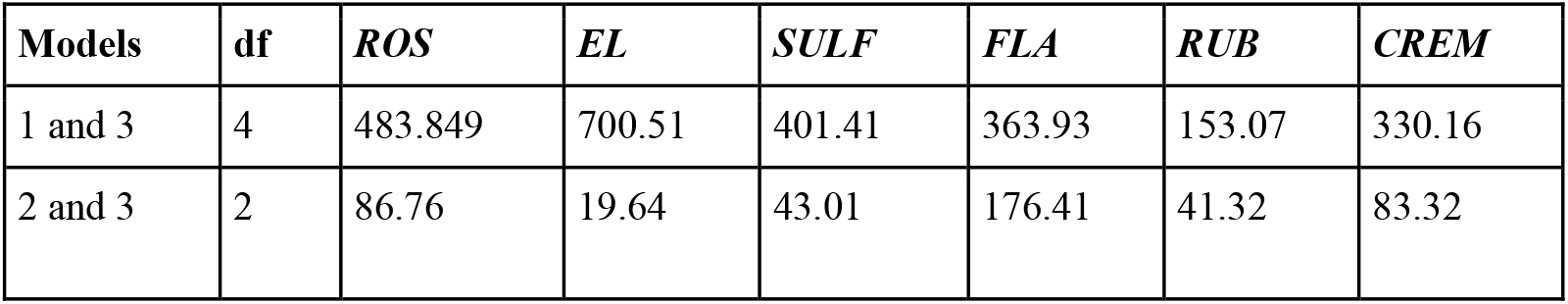
Test statistic from the likelihood ratio test for comparison between models 1 (sigmoid), 2 (sigmoid with flanking polymorphism) and 3 (stepped) cline models. Models 1 and 3 differ by 4 degrees of freedom (df) and models 2 and 3 by 2 degrees of freedom. The critical values associated with the 95% confidence intervals are 9.448 and 5.991 respectively. The more complex model is chosen since the test statistic is greater than the critical value.

**Table S3:**
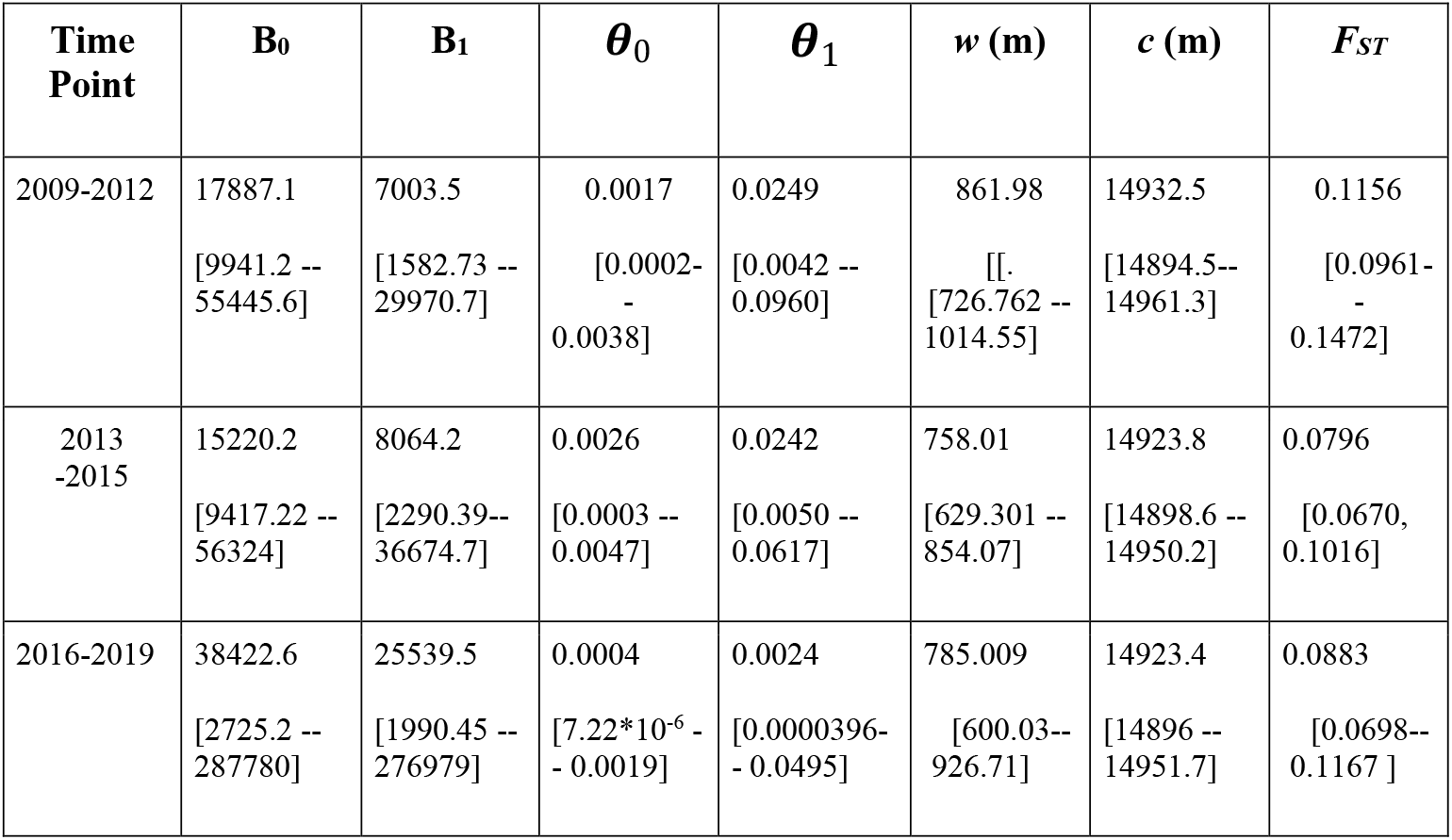
MLE together with the 95% confidence interval (CI) for the parameter from the asymmetric cline model for *ROS* for three time points 2009-2012, 2013-2015 and 2016 to 2019.

**Table S4:** 1D transect divided into different bins (or intervals) with the number of demes (with at least 10 individuals) in each bin and the total number of individuals in all demes.

| Interval (km) | Number of demes | Total number of individuals |
| --- | --- | --- |
| 0 to 12.5 | 9 | 146 |
| 12.5 to 13.25 | 29 | 580 |
| 13.25 to 13.75 | 25 | 621 |
| 13.75 to 14.25 | 28 | 992 |
| 14.25 to 14.5 | 33 | 1420 |
| 14.5 to 14.75 | 29 | 1473 |
| 14.75 to 15 | 37 | 3714 |
| 15 to 15.25 | 39 | 4322 |
| 15.25 to 15.75 | 42 | 5710 |
| 15.75 to 16.25 | 27 | 786 |
| 16.25 to 18.5 | 27 | 632 |
| > 18.5 | 12 | 211 |

**Table S5:** Mean *F*_*IS*_ across demes at each locus compared with the mean from 1000 shuffled replicates (*F*_*IS, null*_) at 25m cluster diameter. The latter represents a null model wherein each deme is at HW equilibrium. The last two columns show the fraction of demes with significantly positive and negative *F*_*IS*_ as compared to the null model. Significance is based on p-value obtained for each deme, which is the fraction of replicates that gives *F*_*IS*_ greater than observed (when *F*_*IS*_ > 0) and less than observed otherwise. Significant heterozygote deficit is observed only at *ROS* (marked by an asterisk) with more than 5% demes with observed *F*_*IS*_ greater than mean *F*_*IS, null*_.

| SNP | Mean $F_{IS}$ | Mean $F_{IS, null}$ | % of demes with significantly positive $F_{IS}$ | % of demes with significantly negative $F_{IS}$ |
| --- | --- | --- | --- | --- |
| *ROS* * | 0.034 | -0.021 | 0.083 | 0 |
| *SULF* | -0.018 | -0.022 | 0.036 | 0.024 |
| *FLA* | -0.016 | -0.021 | 0.032 | 0.019 |
| *RUB* | -0.053 | -0.035 | 0.012 | 0.03 |
| *CREM* | -0.008 | -0.022 | 0.046 | 0.012 |

**Table S6:** Fig S6: *F*_*IS*_ and LD across hybrid zone at 25m cluster diameter: Mean *F*_*IS*_ across 5 unlinked loci and mean correlations 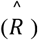 across 10 unlinked locus pairs for different intervals. The table also shows the standard deviations (sd) within each bin, mean and sd of the null distribution for *F*_*IS*_ and LD obtained from 1000 shuffled replicates. P_mean refers to the p-value comparing the means respectively.

| Interval (km) | Mean $F_{IS}$ | sd $F_{IS}$ | Mean $F_{IS, null}$ | sd $F_{IS, null}$ | P_mean | Mean $R$ | sd $R$ | Mean $R_{null}$ | sd $R_{null}$ | P_mean |
| --- | --- | --- | --- | --- | --- | --- | --- | --- | --- | --- |
| 0 to 12.5 | -0.111 | 0.09 | -0.04 | 0.132 | 0.948 | 0.054 | 0.131 | -0.000262 | 0.055 | 0.005 |
| 12.5 to 13.25 | -0.011 | 0.116 | -0.03 | 0.101 | 0.144 | 0.086 | 0.142 | 0.0000291 | 0.065 | 0 |
| 13.25 to 13.75 | 0.015 | 0.168 | -0.028 | 0.097 | 0.01 | 0.128 | 0.193 | 0.0000732 | 0.069 | 0 |
| 13.75 to 14.25 | -0.032 | 0.084 | -0.022 | 0.09 | 0.713 | 0.019 | 0.087 | -0.0002 | 0.058 | 0.044 |
| 14.25 to 14.5 | -0.049 | 0.143 | -0.034 | 0.107 | 0.798 | 0.02 | 0.092 | 0.000136 | 0.080 | 0.089 |
| 14.5 to 14.75 | -0.008 | 0.114 | -0.018 | 0.081 | 0.25 | 0.027 | 0.068 | -0.00062 | 0.061 | 0.013 |
| 14.75 to 15 | -0.024 | 0.116 | -0.028 | 0.093 | 0.353 | 0.049 | 0.076 | 0.00042 | 0.06 | 0 |
| 15 to 15.25 | -0.0002 | 0.090 | -0.016 | 0.079 | 0.092 | 0.029 | 0.074 | -0.00052 | 0.062 | 0.003 |
| 15.25 to 15.75 | 0.020 | 0.087 | -0.015 | 0.075 | 0.003 | 0.017 | 0.059 | 0.00026 | 0.056 | 0.041 |
| 15.75 to 16.25 | -0.014 | 0.104 | -0.024 | 0.098 | 0.303 | 0.02 | 0.082 | -0.000095 | 0.059 | 0.047 |
| 16.25 to 18.5 | -0.032 | 0.143 | -0.03 | 0.112 | 0.526 | 0.052 | 0.133 | -0.00054 | 0.064 | 0 |
| > 18.5 | -0.038 | 0.078 | -0.032 | 0.091 | 0.559 | 0.006 | 0.055 | -0.0004 | 0.045 | 0.304 |

**Table S7:** Mean correlations 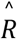 between each pair of loci across demes compared with the mean from 1000 shuffled replicates 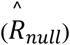 at 25m cluster diameter. Shuffling is done to remove any associations between loci but preserves observed HW deviations. The last two columns show the fraction of demes that show significant positive and negative correlations when compared to the null. All locus pairs has significantly positive observed correlations except for *SULF* and *RUB* and *CREM* and *RUB*, while no locus pairs shows significantly negative correlations.

| SNP 1 | SNP 2 | Mean $\hat{R}$ | Mean $\hat{R}_{null}$<br>(* $10^{-4}$) | % of demes with significantly positive $\hat{R}$ | % of demes with significantly negative $\hat{R}$ |
| --- | --- | --- | --- | --- | --- |
| *ROS* | *SULF* | 0.049 | 2.15 | 0.063 | 0.021 |
| *ROS* | *FLA* | 0.089 | -1.14 | 0.141 | 0.183 |
| *ROS* | *RUB* | 0.029 | 6.33 | 0.082 | 0.034 |
| *ROS* | *CREM* | 0.064 | 1.41 | 0.105 | 0.012 |
| *SULF* | *FLA* | 0.022 | 0.70 | 0.069 | 0.03 |
| *SULF* | *RUB* | -0.002 | -1.83 | 0.045 | 0.048 |
| *SULF* | *CREM* | 0.031 | 2.14 | 0.065 | 0.039 |
| *FLA* | *RUB* | 0.035 | 1.82 | 0.073 | 0.033 |
| *FLA* | *CREM* | 0.066 | -0.74 | 0.121 | 0.027 |
| *CREM* | *RUB* | 0.031 | -1.9 | 0.045 | 0.039 |

**Table S8:** A) Improbable individuals on the yellow flank; there are 27/729=3.7% individuals with P < 2 × 10^−4^ in the yellow flank (defined as < 13km). Probabilities are calculated from the allele frequencies in either flank, and the most likely assignment is shown in bold. Genotypes are roughly classified by eye (13 opposing homozygotes (red); 8 F1-like (italics); 6 foreign homozygotes (bold)). This classification approximately corresponds to the probabilities of coming from the local population, F1, or being a direct seed disperser. However, these probabilities do not show that chance of coming from the centre of the hybrid zone, which is the most likely explanation for the opposing homozygotes. Most individuals come from the immediate flank (between 2.5 and 1.5 km from the centre), but two come from further away. B) The same for the magenta flank, where there are 17/1596=1.06% individuals with P < 2 × 10^−4^ (defined as >16km). Genotypes are roughly classified by eye (8 opposing homozygotes (red); 2 F1-like (italics); 7 foreign homozygotes (bold)). There is a cluster of yellow individuals ∼3.8km out.

| Index | Genotype | z | $\log_{10}(P_0)$ | $\log_{10}(P_{B1})$ | $\log_{10}(P_{F1})$ | $\log_{10}(P_{opp})$ |
| --- | --- | --- | --- | --- | --- | --- |
| S6684 | **{2,2,1,2,0}** | 1646.62 | -8.56024 | -5.01054 | -4.38194 | **-1.83974** |
| P0474 | *{1,1,1,2,1}* | 10722.9 | -3.99329 | -1.9702 | **-1.03414** | -2.77608 |
| M3104 | *{1,1,2,1,1}* | 12046.8 | -4.65039 | -2.82286 | **-2.02388** | -3.21678 |
| J1675 | *{1,2,2,1,1}* | 12812.8 | -5.02378 | -3.18803 | **-2.33861** | -2.75338 |
| M1198 | **{2,0,1,2,0}** | 12817.8 | -3.92564 | **-2.97617** | -3.15966 | -3.36882 |
| M0381 | **{2,2,1,1,2}** | 12820.2 | -6.02382 | -3.94979 | -3.04573 | **-1.49201** |
| M0373 | **{2,1,2,0,2}** | 12830.8 | -7.05989 | -5.14305 | -4.76543 | **-2.99839** |
| P2906 | *{1,1,2,1,0}* | 12832.1 | -3.76651 | -2.68385 | **-2.47669** | -3.96958 |
| M0395 | **{2,2,2,1,1}** | 12832.4 | -6.19668 | -4.10984 | -3.21749 | **-1.52764** |
| M0399 | **{2,2,1,2,1}** | 12834.1 | -5.16352 | -3.17915 | -2.21766 | **-1.08694** |
| L1318 | **{2,1,0,2,1}** | 12834.1 | -3.73422 | -2.61868 | -2.31636 | **-2.26751** |
| S4464 | **{2,1,1,2,0}** | 12835.8 | -3.91184 | -2.67664 | -2.35636 | **-2.30314** |
| M0391 | **{2,2,2,1,0}** | 12836.3 | -5.53804 | -4.0289 | -3.6852 | **-2.28044** |
| M0393 | **{2,1,1,2,0}** | 12836.5 | -3.91161 | -2.67653 | -2.35627 | **-2.30314** |
| P2969 | **{2,1,2,1,0}** | 12840.6 | -4.94267 | -3.60618 | -3.35527 | **-2.74384** |
| M0354 | **{1,0,2,2,0}** | 12843.6 | -4.17692 | **-3.23211** | -3.42623 | -4.47963 |
| M1611 | *{2,2,1,1,1}* | 12855.3 | -4.75078 | -2.92378 | -2.06655 | **-1.64257** |
| S4446 | *{2,1,1,1,1}* | 12880.0 | -4.149 | -2.49915 | **-1.73591** | -2.10597 |
| L1316 | **{1,2,2,1,2}** | 12887.4 | -6.25639 | -4.19772 | -3.3071 | **-2.60282** |
| J0775 | **{2,2,2,2,1}** | 12888.4 | -6.5743 | -4.34767 | -3.35467 | **-0.972009** |
| J0766 | **{2,2,2,1,0}** | 12889.6 | -5.5187 | -4.01895 | -3.67684 | **-2.28044** |
| J0768 | **{2,1,2,0,2}** | 12904.6 | -7.02967 | -5.12815 | -4.75111 | **-2.99839** |
| V6027 | **{1,2,0,1,2}** | 12938.3 | -3.9783 | -2.87661 | **-2.58817** | -3.43492 |
| P2971 | *{1,1,1,1,2}* | 12981.9 | -4.20295 | -2.59056 | **-1.83225** | -3.18115 |
| P0888 | *{2,1,1,1,2}* | 12983.0 | -5.3637 | -3.49766 | -2.69403 | **-1.95541** |
| P2974 | **{1,1,2,2,2}** | 12983.8 | -6.02567 | -4.00816 | -3.11192 | **-2.51059** |
| M0398 | **{0,0,2,2,2}** | 12985.8 | -5.47859 | **-4.43498** | -5.44234 | -5.40259 |

(B)
| Index | Genotype | z | $\log_{10}(P_o)$ | $\log_{10}(P_{B1})$ | $\log_{10}(P_{F1})$ | $\log_{10}(P_{opp})$ |
| --- | --- | --- | --- | --- | --- | --- |
| V6899 | {0,0,0,2,0} | 18375.8 | -9.30623 | -6.11531 | -5.31199 | -1.26318 |
| V6898 | {0,0,0,1,1} | 18375.7 | -9.02308 | -5.62276 | -4.52542 | -1.6172 |
| V6310 | {1,0,0,0,0} | 18375.5 | -8.81046 | -5.21527 | -4.00491 | -1.65773 |
| V6886 | {0,0,0,2,0} | 18375.5 | -9.30577 | -6.11504 | -5.31174 | -1.26318 |
| V6896 | {1,1,1,2,1} | 18375.4 | -3.74179 | -1.48487 | -0.861017 | -3.42887 |
| S8697 | {0,1,0,1,0} | 18374.1 | -8.76883 | -5.36898 | -4.25791 | -0.933579 |
| S0611 | {*,1,0,2,0} | 18205.5 | -3.71579 | -2.85534 | -2.27578 | -1.47056 |
| M1369 | {0,2,1,1,1} | 16797.5 | -4.79714 | -3.21285 | -2.8562 | -3.12103 |
| KTPla8 | {0,1,0,0,0} | 16770. | -7.73874 | -5.17588 | -4.15807 | -1.14172 |
| P0057 | {0,0,0,1,0} | 16679.3 | -7.41817 | -4.93055 | -3.92265 | -0.869115 |
| M0815 | {0,1,1,2,2} | 16412.9 | -4.31431 | -3.26738 | -3.5834 | -4.19875 |
| M0818 | {0,1,0,2,1} | 16407.4 | -4.89528 | -3.33361 | -2.83463 | -2.07574 |
| T4892 | {1,0,0,0,1} | 16358.9 | -5.60791 | -3.67708 | -2.77922 | -2.40582 |
| T3919 | {0,2,0,2,1} | 16320.5 | -4.41297 | -3.27854 | -3.20304 | -2.74243 |
| M0460 | {1,0,2,2,0} | 16302.8 | -4.07101 | -2.99048 | -3.19363 | -3.99123 |
| Z2879 | {1,0,0,*,0} | 16213.4 | -4.63698 | -3.06893 | -2.31586 | -1.59192 |
| S5374 | {0,1,1,1,1} | 16061.3 | -4.27146 | -2.88912 | -2.29752 | -2.45434 |

## Notes

### Competing Interest Statement

The authors have declared no competing interest.

